# *OsPSTOL1* is prevalent in upland rice and its expression in wheat enhances root growth and hastens low phosphate signaling

**DOI:** 10.1101/2022.11.03.515113

**Authors:** Alek Thomas Kettenburg, Miguel Angel Lopez, Kalenahalli Yogendra, Matthew J. Prior, Teresa Rose, Sabrina Bimson, Sigrid Heuer, Stuart John Roy, Julia Bailey-Serres

## Abstract

*PHOSPHORUS-STARVATION TOLERANCE 1* (*OsPSTOL1*) benefits crown root growth and phosphorus (P) sufficiency in rice (*Oryza sativa* L.). To better understand the importance of this variably present gene, we carried out a biogeographic survey of landraces and cultivars, confirming that functional *OsPSTOL1* alleles prevail in low nutrient and drought-prone rainfed ecosystems, whereas loss-of-function alleles and absence haplotypes predominate in control-irrigated paddy varieties of east Asia. To address the evolutionary history of *OsPSTOL1* and related genes in cereal crops, phylogenetic and transcript meta-analyses were performed. Finally, to evaluate its potential value in another Gramineae, wheat (*Triticum aestivum* L.) lines overexpressing *OsPSTOL1* were evaluated under field and controlled low P conditions. *OsPSTOL1* enhances growth, crown root number, and overall root plasticity under low P in wheat. Survey of root and shoot crown transcriptomes at two developmental stages identifies transcription factors that are differentially regulated in *OsPSTOL1* wheat that are similarly controlled by the gene in rice. In wheat, *OsPSTOL1* alters the timing and amplitude of regulators of root development in dry soils and hastens induction of the core P-starvation response. Based on these findings, *OsPSTOL1* and related genes may aid more sustainable cultivation of cereal crops.

**Summary statement:** Might a rice gene that controls root plasticity confer a similar benefit in another grain crop. Here, we evaluate the genetic variation and evolutionary history of *OsPSTOL1* and demonstrate its impact in wheat.

## 1. INTRODUCTION

To feed a growing global population the production of major grain crops must double while simultaneously increasing sustainability (Ray, Mueller, West & Foley 2013; Zhao *et al*. 2017). The essential nutrients nitrogen (N) and phosphorus (P) are critical for optimal growth and yield but their overuse is highly damaging to the environment (López-Arredondo, Leyva-González, González-Morales, López-Bucio & Herrera-Estrella 2014; Yu, Keitel, Zhang, Wangeci & Dijkstra 2022). Although an estimated 18% of global crop land receives insufficient P fertilization, approximately 46% receives an excess that leads to eutrophication of aquatic ecosystems (Carpenter 2008; MacDonald, Bennett, Potter & Ramankutty 2011; Kleinman *et al*. 2015). These environmental challenges, along with the rising cost of P fertilizer and low bioavailability of P in many soils has made the improvement of P use efficiency of crops a critical need (Alewell *et al*. 2020). This might be achieved through introduction of genes that enhance developmental plasticity or metabolic processes that limit loss of quality and yield under P or other nutrient deficiencies.

The rice (*Oryza sativa L*.) *PHOSPHORUS-STARVATION TOLERANCE 1* (*OsPSTOL1*) gene was identified in the upland *aus* landrace Kasalath and found to increase P uptake and yields in P deficient soil by promoting early crown root growth (Gamuyao *et al*. 2012). *OsPSTOL1* lies within a ∼90 Kb insertion-deletion (indel) region on chromosome 12 that is absent from the Nipponbare reference genome (*japonica* subspecies) (Chin *et al*. 2011). Introduction of the Kasalath (K) allele of *OsPSTOL1* into varieties that lack it by breeding or transgenesis confers increased tolerance to P deficiency (Wissuwa & Ae 2001b; Gamuyao *et al*. 2012). However, whether *OsPSTOL1* is an adaptive gene and how it influences root architecture or P sufficiency in rice or when expressed in another monocot crop is an open question.

*OsPSTOL1* is present in *Oryza* species with AA genomes, with most maintaining both gene present and absence alleles (Pariasca-Tanaka *et al*. 2014), suggesting it is under balancing selection (Vigueira, Small & Olsen 2016). Surveys of small collections report that *OsPSTOL1* is present in varieties grown in rainfed upland regions (Chin *et al*. 2010, 2011), where soil nutrient availability is generally lower and drought is prevalent (Haefele, Nelson & Hijmans 2014). *OsPSTOL1* is part of an uncharacterized family of putative kinases with homologs in other Gramineae, including *Zea mays* (maize) (Azevedo *et al*. 2015), *Triticum aestivum* (wheat) (Milner *et al*. 2018), and *Sorghum bicolor* (sorghum) (Hufnagel *et al*. 2014). *OsPSTOL1* homologs in maize and sorghum colocalize with quantitative trait loci for root morphology and P uptake (Hufnagel *et al*. 2014; Azevedo *et al*. 2015), suggesting that functions of *OsPSTOL1*-like genes may be conserved; yet their origins and distinctions in structure and expression remain unclear.

Here, we survey *OsPSTOL1* presence/absence and allelic variation across more than 3000 re-sequenced landrace and varietal genomes and assess the biogeographical and dispersal history of these alleles (Gutaker *et al*. 2020). We find that the characterized K allele of *OsPSTOL1* is uncommon in paddy and control-irrigated rice and overrepresented in accessions of rainfed ecosystems that are prone to drought. Wheat is also a predominantly rainfed crop, but its genes underlying P uptake and root architecture are understudied (Oono *et al*. 2013b; Walkowiak *et al*. 2020). To explore translatability of the adaptive function of *OsPSTOL1*, we examine the impact of its ectopic expression in wheat and identify increases in growth under P-replete and limiting conditions in multiple environments, including the field. Evaluation of *OsPSTOL1* effects on roots and crowns demonstrates that *OsPSTOL1* influences root architecture, yield traits and transcripts of regulators of developmental plasticity that are important under drought and in low P sensing.

## 2. MATERIALS AND METHODS

### 2.1 Determination of genotypic, geographic and cultivation ecosystem information of rice varieties

Haplotype information of *OsPSTOL1* and neighboring loci were determined using vcf files and fastq files obtained from the 3000 rice genome project (Wang *et al*. 2018b), Rice Pan-Genome Browser (Sun *et al*. 2017) and Gutaker *et al*. (Gutaker *et al*. 2020). Country-of-origin, subspecies, rice growing ecosystem, and SNP group information was mined from the Rice Pan-Genome Browser (Sun *et al*. 2017) and (Gutaker *et al*. 2020). Countries with fewer than 10 varieties sampled were discounted. Regional data on rice growing ecosystem distribution were from (Haefele *et al*. 2014). P fertilizer application data were from (Roser & Ritchie 2013), updated in 2017. See Supporting Information: Methods for details of genome and allele evaluation.

### 2.2 *OsPSTOL1* homologs, phylogenetic tree construction, and protein domain prediction

*OsPSTOL1* orthologs have been reported in some *Oryza* (Pariasca-Tanaka *et al*. 2014; Neelam *et al*. 2017). To identify *OsPSTOL1*-like genes in other species, BLASTP was performed with the K-allele of OsPSTOL1 as a query against proteomes of related species. Gaps were trimmed with trimAl (Capella-Gutiérrez, Silla-Martínez & Gabaldón 2009) using default settings. Tree was constructed using RAxML (Kozlov, Darriba, Flouri, Morel & Stamatakis 2019), with default settings and 1000 bootstraps. Protein domains were predicted using hmmscan (Finn, Clements & Eddy 2011), ScanProsite (de Castro *et al*. 2006), and Phobius (Käll, Krogh & Sonnhammer 2007). See Supporting Information: Methods for more detail.

### 2.3 Generation of *pUbi::OsPSTOL1* wheat lines

The coding sequence of *OsPSTOL1* (*OsPUPK46-2*; BAK26566.1) of Kasalath was synthesized, ligated into *pENTR/D-TOPO* (Invitrogen, Carlsbad, CA, USA), validated by sequencing, and transferred by Gateway LR recombination (Invitrogen) into the T-DNA vector *pMDC99* (Curtis & Grossniklaus 2003), which contains a *Zea mays Ubiquitin1* (*Ubi1*) promoter (JX947345.1) for driving genes expression and a *35S CaMV* terminator. *Agrobacterium*-mediated transformation of wheat, cv. Fielder, with *Ubi::OsPSTOL1* was conducted (Ishida, Tsunashima, Hiei & Komari 2015). Three independent *OsPSTOL1* lines (T_3_-T_5_) were compared to a null segregant from the T_1_ generation. See Supporting Information: Methods for detail on transgene evaluation. Plasmids and wheat genetic material are available from S.R.

### 2.4 Evaluation of wheat under greenhouse and field conditions

Evenly sized seeds were imbibed for 4 h and placed in the dark at 4 ºC for 3 d prior to transplanting into pots (diameter 150 mm and height of 150 mm) filled with a 2.0 L of cocopeat soil (South Australian Research and Development Institute, Adelaide, Australia). Plants were grown in The Plant Accelerator (Australian Plant Phenomics Facility, University of Adelaide, Australia (−34.971353 S; 138.639933 E) with natural light and 22ºC daytime and 15ºC night temperatures. Grain protein content and total grain nitrogen was measured using a nitrogen analyzer (Rapid N exceed®, Elementar, Germany). Chlorophyll content was measured using a SPAD-502Plus (Konica Minolata Inc, Japan).

Field evaluation was at Glenthorne Farm, South Australia (−35.056923 S 138.556224 E) between May and December 2018. T_5_ null segregant seeds were used as the control. The fertilized plot (50 g m^2^ High Phosphate Super Phosphate (RICHGRO, Jandakot, Western Australia) and 20 g m^2^ Soluble Nitrogen Urea (RICHGRO)) was planted and managed and evaluated weekly as described previously (Regmi *et al*. 2020), with irrigation by rainfall. Blitzem Snail & Slug Pellets (Yates, Auckland, New Zealand) was applied at a rate of 5 h m^2^ to avoid issues with snails. Biomass was measured as the average biomass of all plants per genotype within a plot (two plots per genotype). For grain nutrient analysis, plants were divided into four groups of eight, and the grain from each group was pooled as one biological replicate and nutrients measured using microwave digestion followed by ICP-OES.

### 2.5 Evaluation of wheat under controlled P nutrition

Seedlings of average size and growth were transplanted into each pot containing a pre-washed mix of 70% (v/v) #20 silica sand (Gillibrand) and 30% (v/v) Green Grades Profile™. Profile™ contains 10% (v/v) aluminum oxide, which acts as a solid-state buffer for P (Lynch, Epstein, Lauchli & Weight 1990). Prewashing with water was performed to reduce the starting P content to 25 μM. Pots were sub-irrigated with 3 L of industrial water supplemented with the additional nutrients. For high [Pi] (HP), the solution contained 436 μM KH_2_PO_4_, and for low [Pi] (LP) this was substituted with 440 μM KCl (Table S1c). Plants were harvested at 36 and 50 DAS, representing Zadoks stage 23 and stage 31, respectively. Each pot (5-6 plants) was pooled as a biological replicate, with four replicates per treatment and sampling time. For evaluation of root systems, pre-germinated seeds were transplanted into 40 cm tall, 10.16 cm diameter PVC pipes filled with the prewashed mix of 70% (v/v) sand and 30% (v/v) Profile™ and drip irrigated daily. For HP, the solution contained 436 μM KH_2_PO_4_ and for LP, 5 μM KH_2_PO_4_ and 335 μM KCl. At 28 DAS, root systems were recovered. See Supporting Information: Methods for more detail. For nutrients, Pi concentration of approximately 150 mg fresh tissue was extracted (Yamaji *et al*. 2017) and measured by the molybdate blue colorimetric method (Murphy & Riley 1962). Total P of 15 to 25 mg of dry tissue was determined by persulfate digestion using a 10 mL reaction of 5% (v/v) sulfuric acid and 6% (w/v) potassium persulfate (Jeffries, Dieken & Jones 1979). Nitrogen (N) in dry tissue was measured using a Flash EA1112 N analyzer at the UC Riverside Environmental Sciences Research Laboratory.

### 2.6 mRNA-seq library construction, sequencing, read processing, and analysis

Three biological replicates were used. TRizol (Invitrogen) extracted total RNA was processed and used to generate libraries for mRNA-sequencing as described previously (Reynoso *et al*. 2019). Sequences were aligned using hisat2 (Kim, Langmead & Salzberg 2015). For wheat, IWGSC v1.1 with the *OsPSTOL1* K-allele CDS added as a separate chromosome. For publicly available RNA-seq data, reads were aligned to these genomes: *Oryza sativa* IRGSP-1, *O. rufipogon* OR_W1943, or v1.0 *O longistaminata*. For mapping transcripts to *OsPSTOL1* alleles, the region from 5000 bp upstream to 500 bp downstream of the start and stop codons was added as a separate chromosome [GenBank AB458444.1, K-allele, N22; GenBank SAMN04568482, representative J-allele; *O. rufipogon* W0141 for *OrPSTOL1* (Zhao *et al*. 2018); *O. longistaminata* KN540838.1_FG002 scaffold KN540838.1: 24,114-24,980 for *OlPSTOL1*]. Alignments were visualized using the Integrated Genome Viewer browser v2.7 (Narang *et al*. 2010). Differentially regulated gene (DRG) transcripts were identified by use of the limma-voom package (Law, Chen, Shi & Smyth 2014) with rawread counts normalized with voom by the quantile method. Genes with less than 5 counts per million (CPM) in at least three samples were excluded. Log_2_ Fold Change (Log_2_FC), adjusted P values (adj.P.Val) and false discovery rate (FDR) were calculated. Clustering was performed (Abu-Jamous & Kelly 2018) using Z-score normalization of Log_2_FC values, followed by Gene Ontology (GO) (H Backman & Girke 2016) using the wheat GO definitions from BioMart (Smedley *et al*. 2009) (Ensembl Release 50). Transcription factors were subsetted using the Plant Transcription Factor Database (Jin *et al*. 2017). Wheat orthologs of rice genes were found using BioMart. See Supporting Information: Methods for details. All sequence read data are available at NCBI Gene Expression Omnibus, https://www.ncbi.nlm.nih.gov/geo/ under accession GSE185875.

### 2.7 Statistical analyses and visualization

The stats package of R (R Core Team 2017) was used for ANOVA, linear modeling, and Chi-square Goodness of fit testing. Graphs were made with ggplot2 (Wickham 2016), ggradar (Bion 2019), and UpSet packages (Lex, Gehlenborg, Strobelt, Vuillemot & Pfister 2014).

## 3 RESULTS

### 3.1 *OsPSTOL1* was lost in a subsection of rice varieties after the temperate-tropical split in *japonica*

A systematic comparison of *OsPSTOL1* alleles of the genotypes represented in the 3000 Rice Genome collection (Sun *et al*. 2017) identified two sub-groups (Figure 1a-c; Table S2a-d). One group has the characterized *OsPSTOL1 K-*allele of *O. sativa* Kasalath and includes accessions of widely distributed wild species including *O. rufipogon* (Asia), *O. glumaepatula* (South America) and *O. longistaminata* (west Africa). The second group has the uncharacterized *O. sativa* J-alleles that are also widely distributed in *O. barthii* (Africa), *O. glaberrima* (Africa), and *O. meridionalis* (Australia) (Pariasca-Tanaka *et al*. 2014; Vigueira *et al*. 2016). The widely dispersed extant descendant of the progenitor of domesticated rice, *O. rufipogon*, has both J- and K-alleles. In addition, our analysis affirms that *OsPSTOL1* was disrupted on multiple occasions by loss-of-function mutations or interstitial chromosomal deletion. In addition to the characterized deletion of the gene in Nipponbare (Heuer *et al*. 2009) (henceforth the Absent haplotype), we find two K-allele variants (henceforth KStop1 and KStop2) (Figure 1b,c; Figure S1). Notably, we found an allele of the *OsPSTOL1* ortholog of *O. meridionalis* with a nonsense mutation that prematurely terminates the open reading frame before the kinase domain. In summary, both domesticated *O. sativa* and broadly dispersed wild *Oryza* posess J- and K-alleles, allele variants resulting in truncation, and absent haplotypes (Table S2b-c) (Pariasca-Tanaka *et al*. 2014; Neelam *et al*. 2017). It is not known if the J- and K-alleles are distinct in terms of gene regulation or protein function, but a survey of transcriptomes from a panel of 38 diverse varieties (Lou *et al*. 2017) finds K-allele genotypes maintain statistically higher transcript abundance in both shallow and deep roots (Figure 1d).

**Figure 1.**
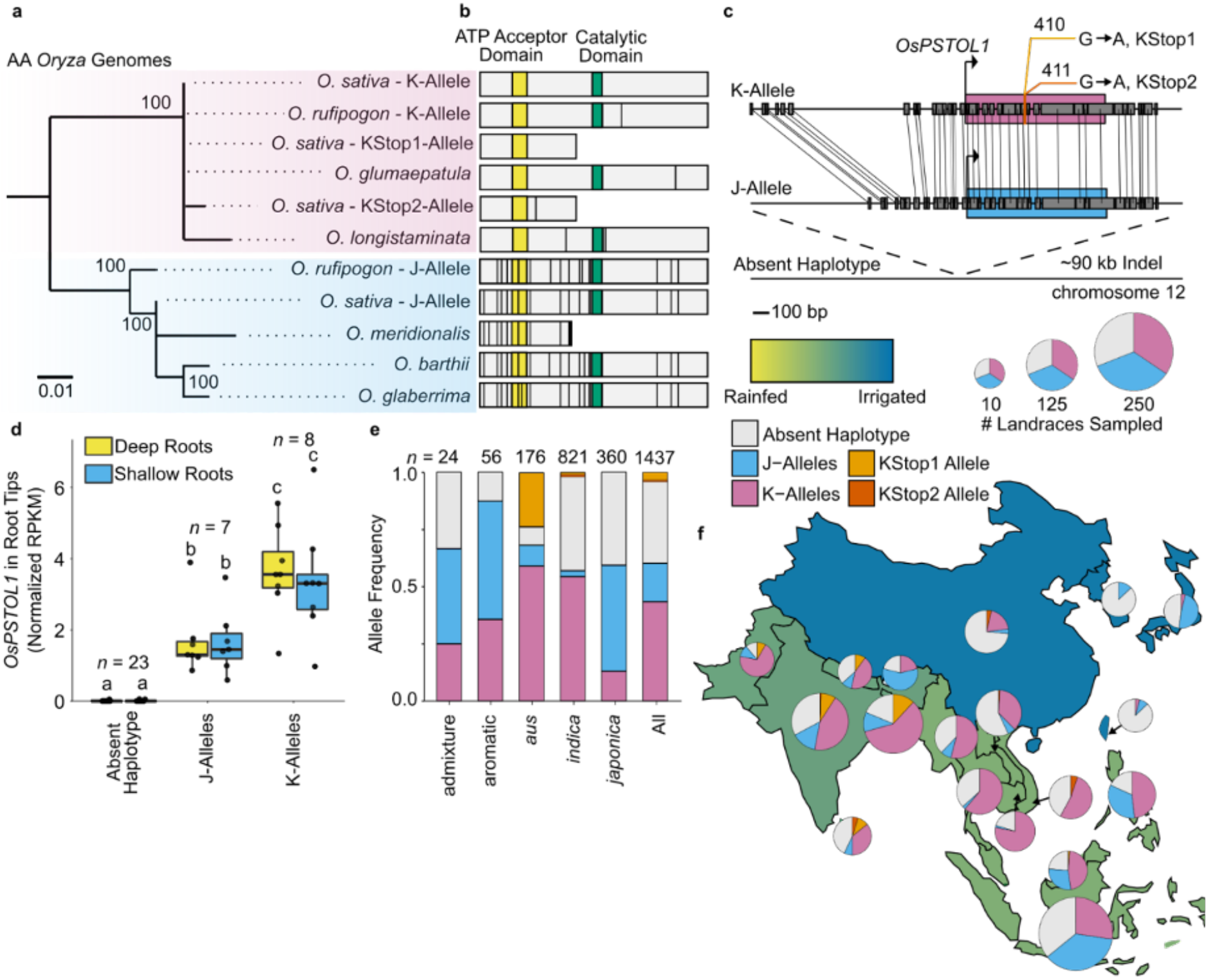
*OsPSTOL1* Haplotype variation and geographic distribution. **a**, Phylogenetic tree based on PSTOL1 amino acid sequences within AA genome *Oryza*. The K-allele group is highlighted magenta, while the J-allele group is highlighted blue. Scale bar indicates amino acid substitutions per position. Node labels indicate the percentage of 1000 bootstraps. **b**, Predicated protein models of PSTOL1 proteins within *Oryza*. Vertical black lines indicate a non-synonymous mutation relative to the K-allele of *O. sativa* (Kasalath landrace). **c**, Genome alignment of *OsPSTOL1* haplotypes in *O. sativa*. Black rectangles indicate segments of common sequence. Colored rectangles represent coding sequences. Arrow facing right indicates the start of translation. The sequence from Kasalath was used for the K-allele, that from N22 for the J-allele and that from Nipponbare for the Absent haplotype. **d**, Abundance of *OsPSTOL1* mRNA in root tips of shallow and deep roots from a diversity panel of rice varieties (Lou *et al*. 2017). Letters indicate significant differences (ANOVA followed by Tukey’s HSD, *P*<0.05). Boxplot boundaries represent the first and third quartiles; a horizontal line divides the interquartile range, median **e**, Distribution of *OsPSTOL1* haplotypes in landraces by subspecies. **f**, Geographic distribution of *OsPSTOL1* haplotypes in landraces. Pie charts are scaled to the number of landraces sampled from each country. Color of each region indicates the proportion of cultivated rice land that is rainfed.

The similar broad species distribution of *PSTOL1* presence and Absence haplotypes within *Oryza* species indicates selection for variants with high transcript accumulation as well as non-functionality including gene loss. This trans-*Oryza* genetic variation could reflect adaptive or undesirable roles in particular environments. To begin to address this, we considered *OsPSTOL1* variation based on geographic and collection attributes of 1438 *O. sativa* landraces (Wang *et al*. 2018b; Gutaker *et al*. 2020). A total of 64% of the landraces have *OsPSTOL1*: functional J-alleles predominate in *japonica* and aromatic landraces (17%) and K-alleles predominate in *aus* and *indica* landraces (43%) (Figure 1e). Landraces with the absent *OsPSTOL1* haplotype predominate in East Asia (Figure 1f).

To evaluate the provenance of *OsPSTOL1* variation, we leveraged the reconstruction of the historic distribution of landraces across Asia generated by Gutaker *et al*. (2020), who identified eight and six modern geographic subpopulations of *japonica* and *indica* rice, respectively (Figure 2a,b). *OsPSTOL1* presence is near ubiquitous in the genetically distinct *japonica* subpopulations, except in landraces of Laos and Taiwan/Japan (Figure 2a,c,d). Of the temperate *japonica*, the Taiwan/Japan population consists almost entirely of lowland irrigated landraces, whereas a J-allele group, containing Korea/Japan landraces, is predominantly grown in upland farms, where soils are more aerobic than irrigated lowland paddies (Kirk, Greenway, Atwell, Ismail & Colmer 2014). Based on the landrace dispersal pattern, *OsPSTOL1* was lost in the temperate Taiwan/Japan lineage after the temperate-tropical split of the *japonicas*, an estimated 4100 yr BP (Figure 2d).

**Figure 2.**
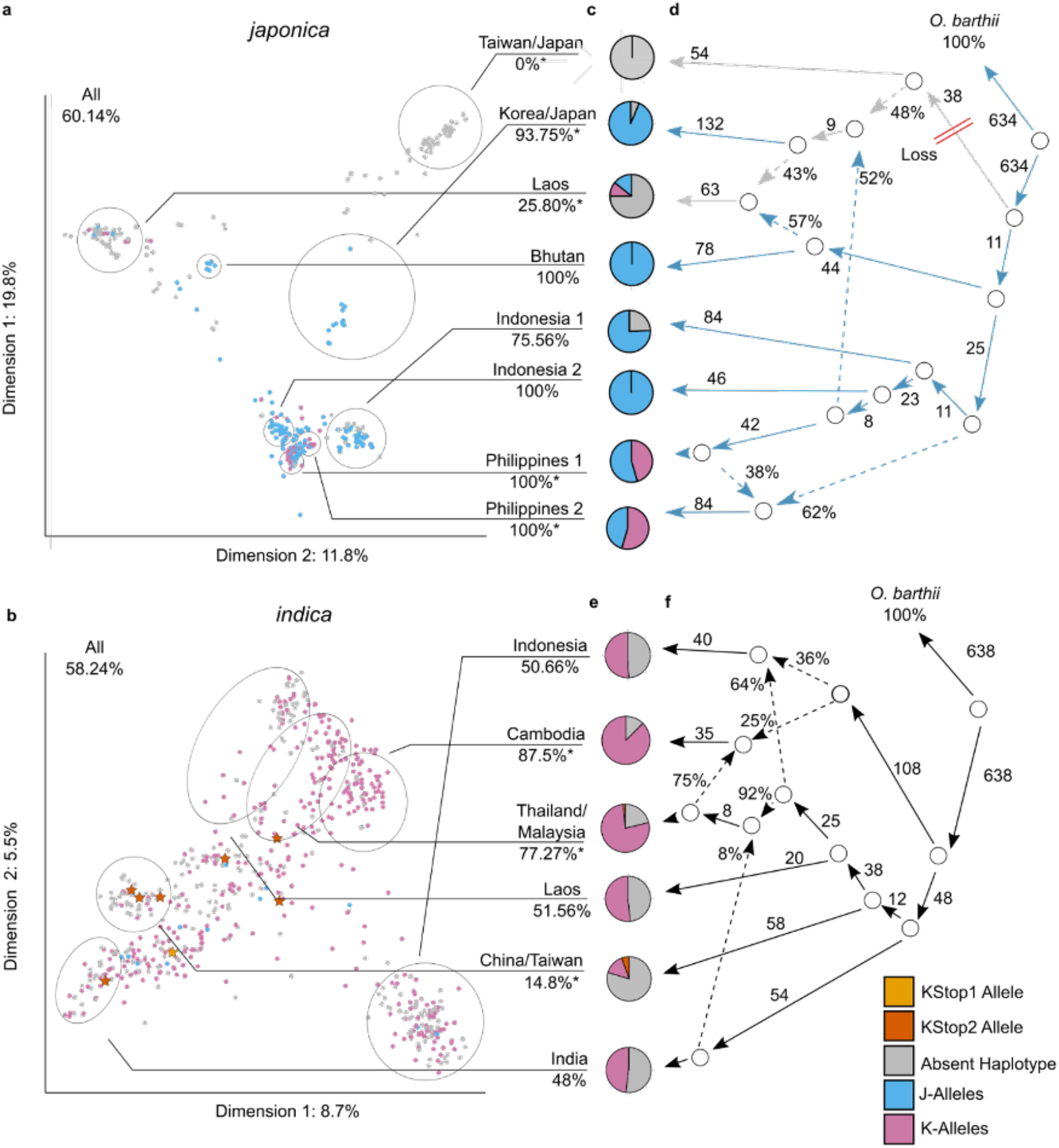
Distribution of *OsPSTOL1* in discrete geographic subpopulations of rice landraces. *OsPSTOL1* SNP variation data are superimposed onto a multidimensional scaling analysis by geographic region and deduced ancestries of 225 *japonica* and 690 *indica* landraces exactly as reported by Gutaker et al. (2020). **a,b**, *japonica* (**a**) and *indica* (**b**) landraces projected onto the first two dimensions after multidimensional scaling of genomic distance data. Colors indicate *OsPSTOL1* alleles. KStop1- and KStop2-alleles are indicated as stars, whereas other alleles as circles. **a**, Gutaker et al. (2020) clustered *japonica* genotypes using *k*-medoids (*k*= 9 subpopulations) and filtered using silhouette parameters, which resulted in k_d_= 8 discrete subpopulations, contained within grey circles. Taiwan/Japan accessions are cultivated under irrigated lowland conditions. Subpopulation names of correspond to predominant geographic location of individual accessions in each subpopulation. **b**, Gutaker et al. (2020) clustered the *indica* genotypes using k-medoids (*k*=7 subpopulations) and filtered resulting in *k*_d_= 6 discrete subpopulations contained within grey circles. **c,e**, Pie charts representing the allelic composition of each discrete subpopulation of *japonica* (**c**) and *indica* (**e**) subgroups. Percentages indicate the frequency of full J- and K-alleles within each population. An asterisk indicates the frequency of *OsPSTOL1* presence alleles is significantly different than expected by *Chi*-square test (*P*<0.05). **d**, (Gutaker *et al*. 2020) admixture graph for the *k*= 9, *k*_d_= 8 *japonica* subpopulations, rooted with *Oryza barthii* as an outgroup. *PSTOL1* is found in all *O. barthii* varieties surveyed (Pariasca-Tanaka *et al*. 2014). Graph represents topology that is consistent between models for all lower values of *k*. Two red slashes indicate the predicted loss of *OsPSTOL1* in the temperate lineage. Blue arrows indicate predicted routes of dispersal of the J-allele group, while grey arrows represent predicted routes of dispersal of the Absent haplotype. **f**, (Gutaker *et al*. 2020) admixture graph for *k*= 7, *k*_d_= 6 indica subpopulations of **c**, rooted with *O. barthii* as an outgroup. **d,f**, Solid arrows represent the predicted uniform ancestries, associated values are estimated scaled drift (*f*_2_) and dashed lines represent mixed ancestries, with estimated proportion of ancestry as a percent.

To clarify the origin of J-alleles in the Korea/Japan population, we tracked dispersal at the haplotype level by following variation at the flanking loci: *OsPUPK20-2* (*LOC_Os12g26380*) and *OsPUPK67-1* (*LOC_Os12g26470*) (Figure 2g; Figure S2a-c; Table S2f). Using a lower-case letter to designate the alleles of these genes, the J-allele of Korea/Japan and Philippines are flanked by *PUPK20-2-m* and *PUPK67-1-a*, whereas those of Taiwan/Japan are bordered by the more distant *PUPK20-2-*e and *PUPK67-1-h*. This determines that the Korea/Japan J-alleles were introgressed from a tropical population that is related to the extant Philippines 2 population (Figure S2a-c), possibly through selection. Our survey also clarifies that K-alleles in *japonica* were introgressed from *indica* landraces, indicating unaccounted admixing (Figure S2a-d). All *japonica* landraces with the Absent haplotype have the *PUPK20-2 a* or *e* allele as well as *PUPK67-1-h*, characteristic of the temperate subpopulation (Figure S2a-d). This confirms they originate from a single deletion event. Reflective of the high levels of admixing in *indica* accessions documented by Gutaker et al. (2020), we find not clear patterns of dispersal of *OsPSTOL1* in the *indica* subspecies, although the more highly expressed K-allele prevails in all but the China/Taiwan subpopulation (Figure 2b,e,f).

### 3.2 *OsPSTOL1* gene presence prevails in rainfed growing systems lacking controlled irrigation

The loss of *OsPSTOL1* in the irrigated Taiwan/Japan *japonicas* and restoration by introgression in upland rainfed landraces of Korea/Japan suggests that this gene may be advantageous in specific cultivation ecosystems. To begin to address this we considered *OsPSTOL1* distribution in landraces known to be cultivated in irrigated lowland, rainfed lowland or uplands, the latter two more prone to drought. This identifies significant enrichment of J-alleles in *japonica* landraces from the rainfed uplands (*P*<0.05) and depletion in irrigated accessions (*P*<0.05) (Figure S3a). In *indica*, K-alleles are depleted in the irrigated lowlands (*P*<0.05) but have similar frequency in the rainfed lowlands and uplands.

To investigate this hypothesis further, we examined the 3000 Rice Genomes that include landraces and varieties of Asian cultivated rice and considered country-based cultivation environment data. Consistent with the biogeographical survey of landraces, the lowest frequency of *OsPSTOL1* (5.92%) is found in the irrigated paddy landraces of Taiwan/Japan *japonica* and their descendants (Single nucleotide polymorphism group JG7) (Figure S3b; Table S2b). The KStop2 allele is nearly exclusive to *indica* in China, consistent with the dispersal of *indica* landraces (Figure 3a,b; Figure S3c). Cambodia, where 88% of the rice is rainfed (FAO 2011a) and droughts are common (Chhinh & Millington 2015), has the highest *OsPSTOL1* and K-allele presence within Asia (Figure 3a,b). By contrast, neighboring Vietnam, which has the highest rate of irrigated rice in the region (FAO 2011b), has low *OsPSTOL1* presence (Figure 3a,b). Laos is exceptional, with the lowest *OsPSTOL1* presence in southeast Asia, despite cultivation of a similar percentage of irrigated rice as Cambodia (FAO 2011c). Outside of Asia, functional *OsPSTOL1* alleles predominate in west Africa and Madagascar, where rice is predominantly rainfed and soil nutrients are generally poor (Figure 3 a,b; Figure S3c) (Haefele *et al*. 2014). Yet, presence and Absence alleles are found in varieties on other continents where irrigation and fertilization are well controlled. We do not know if this distribution reflects selection or ancient and modern dispersal routes of rice from Asia to Europe (Spengler *et al*. 2021) and Asia and Africa to the Americas.

**Figure 3.**
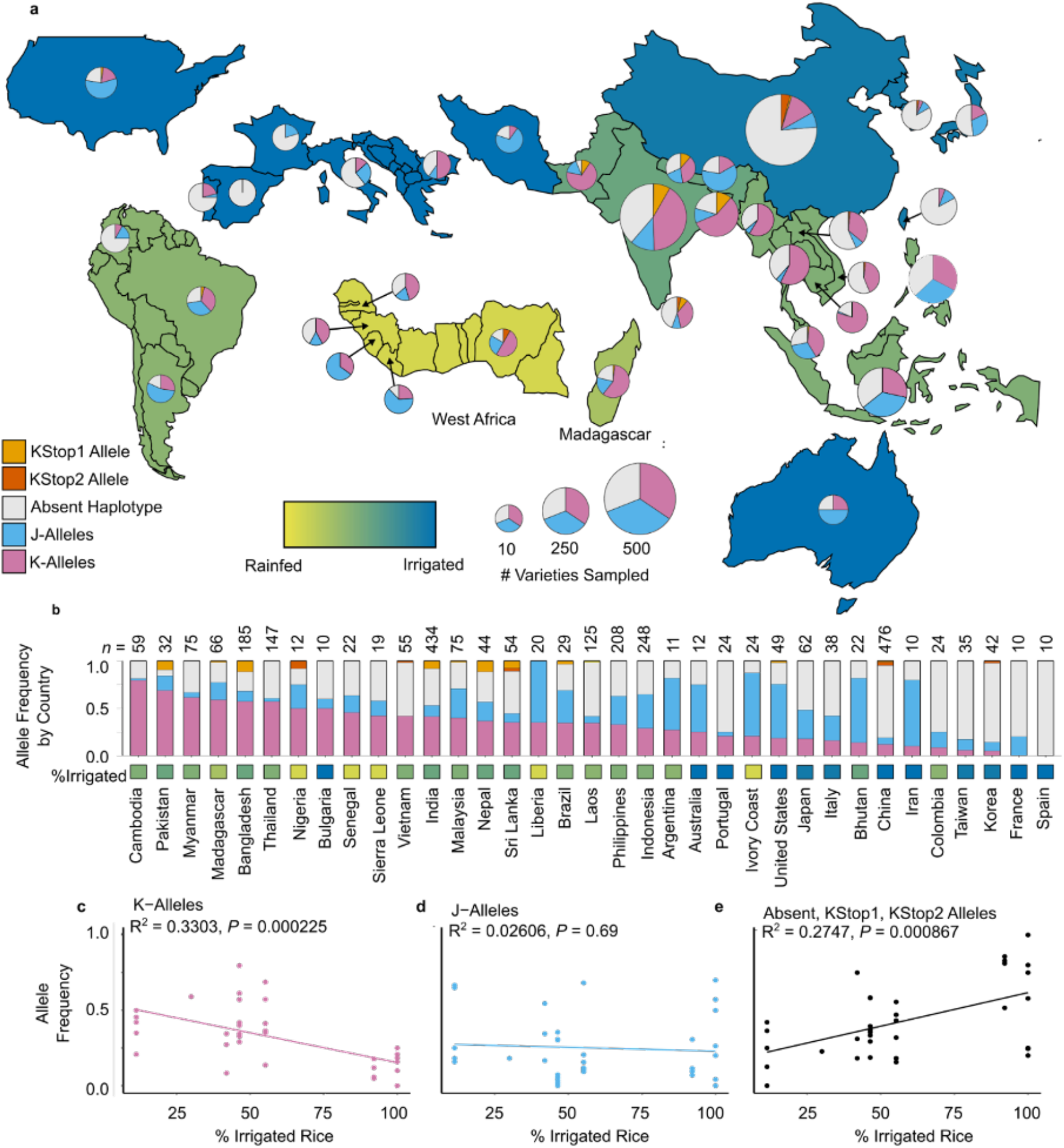
Geographic distribution of *OsPSTOL1* in the 3000 Rice Genomes dataset. **a,b** Distribution of *OsPSTOL1* haplotypes based on the country where the variety was collected. Size of pie charts is proportional to the number of varieties sampled. Color of regions indicates the proportion of cultivated rice land that is irrigated relative to rainfed (lowland or upland). **c-e**, Correlation between the frequency of K-alleles (**c**), J-alleles (**d**), and Absent haplotype (**e**), and the proportion of irrigated rice in a country. R^2^, *P*-value, and best-fit line calculated using linear regression.

Given the somewhat mixed results in terms of cultivation ecosystem and genetic variation, we performed a regression analysis to quantitatively assess allele bias in rainfed versus control irrigated cultivation ecosystems. This determines that global frequency of functional K-alleles negatively correlates with the proportion of irrigated rice (*P*=0.00023), J-alleles show no correlation (*P*=0.69), and the deletion/loss-of-function variants (Absent, KStop1, and KStop2) show a positive correlation with irrigation (*P*=0.00087) (Figure 3c-e). Notably, the frequency of deletion/loss-of-function also positively correlates with P fertilizer application rate (*P*=0.038) (Figure S4c), whereas J-alleles show a weak negative correlation and K-alleles show no correlation (Figure S4a,b).

### 3.3 *OsPSTOL1-like* genes have protein domains characteristic of extracellular signal recognition kinases, yet *OsPSTOL1* and several other family members lack these domains

To gain insight into the evolutionary history of *OsPSTOL1*, the putative *Os*PSTOL1 kinase domain (∼31 kDa) was used to identify *OsPSTOL1*-like proteins in other Gramineae crops. A combined phylogenetic and protein domain analysis of 31 *OsPSTOL1*-like genes identifies rice *OsPSTOL1* and *LOC_Os8g24630*, wheat *TaPSTOL1* (*TraesCS5A02G067800LC*) (Milner *et al*. 2018) and maize *Zm00001d039923* as family members limited to a kinase domain (Figure 4a,b). All other OsPSTOL1-like proteins (36-101 kDa) have N-terminal domains that bind or remodel extracellular carbohydrates (*i.e*., wall-associated kinase (WAK), galacturonan-binding (GUB) WAK (Kanneganti & Gupta 2008; Kohorn & Kohorn 2012), LysM, glycoside hydrolase 18 (GH18) legume lectin domains involved in recognition of carbohydrates from microbes (Mesnage *et al*. 2014; Lannoo & Van Damme 2014; Wang *et al*. 2019). Consistent with their similarity to kinases that recognizes pathogen associated molecular patterns and activate intracellular signaling (Zipfel 2014; Cayrol, Delteil, Gobbato, Kroj & Morel 2016), these genes are induced by microbes and their effectors (Figure S5a-d).

**Figure 4.**
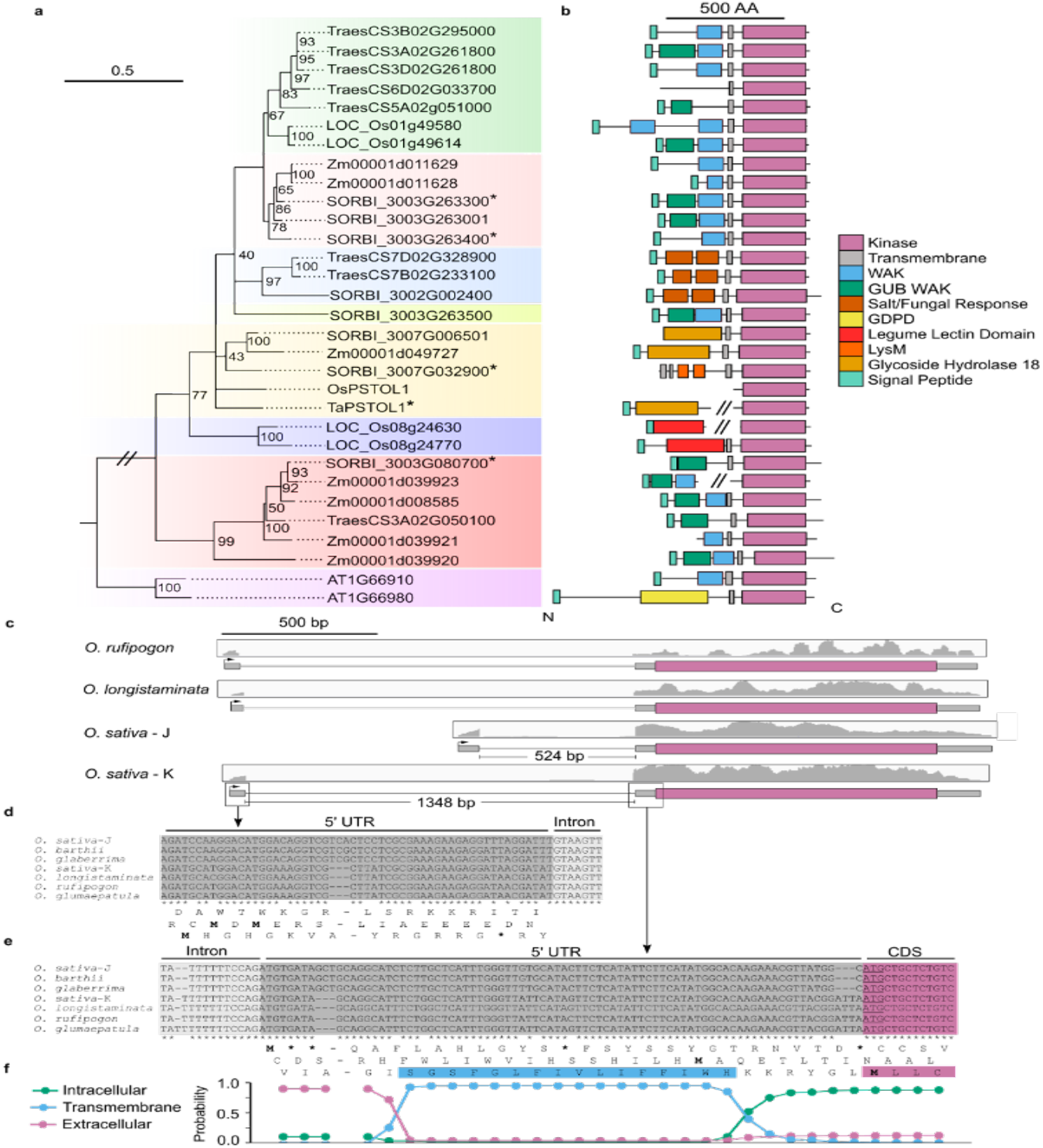
*OsPSTOL1* has an untranslated region encodes a transmembrane domain that is within *OsPSTOL1*-like genes. **a**, Phylogenetic tree of *OsPSTOL1*-like protein in rice *(Oryza sativa, Os*), sorghum (*Sorghum bicolor, SORBI*), maize *(Zea mays, Zm)*, bread wheat *(Triticum aestivum, Traes*), and *Arabidopsis thaliana (At)*. Alignment was based on the amino acid sequence of the kinase domain. For genes with multiple transcript isoforms, the longest deduced protein model was used. Node labels indicate the percentage of 1000 bootstraps. *TaPSTOL1* (TraesCS5A02G067800LC) was formerly annotated as Traes_5AS_AA3DC6A5F. *OsPSTOL1* K-allele used (BAK26566.1). Scale bar indicates amino acid substitutions per position. The tree was constructed by the maximum likelihood method. *SORBI_3003G080700, SORBI_3003G263300, SORBI_3003G263400*, and *SORBI_007G032900* were formerly annotated as *Sb03g006765, Sb03g0031680, Sb03g031690*, and *Sb07g002840*, respectively (Hufnagel *et al*. 2014; Bernardino *et al*. 2019). Genes marked with an asterisk are known to influence growth or development. **b**, Protein models of *OsPSTOL1*-like proteins. Two diagonal lines between two separate segments indicate the aligned kinase domain was downstream of an annotated gene that has high sequence identity to genes near it on the phylogenetic tree. WAK, wall-associated receptor kinase; GUB WAK, wall associated receptor kinase galacturonan-binding; GDPD, glycerolphosphodiesterase. **c**, Alignment of RNA-seq reads from publicly available datasets to the K-allele of *OsPSTOL1* locus in *O. rufipogon* and *O. longistaminata*, and the J- and K-alleles of *O. sativa*. mRNA-seq derived from root tissue was used for *O. rufipogon, O. longistaminata* and the K-allele of *O. sativa*. RNA-seq derived from the crown was used for the J-allele group of *O. sativa*. Gene structure illustrated as a magenta box for the predicted coding sequence, gray box indicates the predicted 5’ untranslated region (UTR), and a black horizontal line for an intron deduced from RNA-sequencing reads are supported by canonical intron splice donor and acceptor sites. RNA sequence reads were visualized with a genome browser and reads mapped in grey within a black horizontal box, placed above the gene structure diagram. **d,e**, Nucleotide sequence alignment of exon 1 and the 5’ end of the intron positioned within the 5’ UTR (**b**) and the 3’ end of the intron and 5’ end of exon 2 that precedes the start codon (**c**) of *OsPSTOL1*. Translation of the three reading frames for *O. sativa* (Kasalath) is shown. **\*** indicates stop codon. Blue highlight indicates the putative transmembrane domain sequence now within the 5’ UTR of the gene. Magenta highlight indicates the start of the translated PSTOL1 protein. **f**, Transmembrane domain prediction calculated using Phobius for the amino acid sequence in the third frame of (**e**). *OsPSTOL1* mRNA translation is predicted to initiate downstream of this domain, given the absence of any 5’ AUG or alternative start codon. See Table S1a for details of the datasets and accessions surveyed.

Next, we considered whether the kinase-only PSTOL1-like proteins arose from multi-module proteins and found evidence in *Triticum* and *Oryza* species. A gene encoding a GH18 domain lies directly upstream of *TaPSTOL1* (Milner et al. 2018), whereas GH18 and kinase domains are present in a single *OsPSTOL1-like* gene (*TRIUR3_01194*) in *Triticum urartu*, the AA genome progenitor of bread wheat (Figure S6a,b). The alignment of mRNA-seq reads to the *PSTOL1* region of *O. sativa, O. rufipogon* and *O. longistaminata* led us to a 5’ exon and intron of variable length that lies upstream of the previously annotated start codon (Figure 4c-f; Figure S6c-f). The 5’ region of the *PSTOL1* gene of all three species has high sequence identity to a transmembrane domain that is in-frame with the kinase domain. However, the transmembrane domain lacks an AUG initiation codon. We conclude that *OsPSTOL1* arose from gene encoding a membrane-anchored kinase. The cellular functions of OsPSTOL1 and related solo-kinase-domain proteins found in other species remain to be determined.

#### *OsPSTOL1* and some related Gramineae genes are associated with plant architecture and nutrition but show mRNA accumulation in varied tissues

Meta-analysis of mRNA-seq data of Gramineae crops reveals that *OsPSTOL1-like* gene transcripts accumulate in diverse tissues, are induced by pathogens and PAMPs, but in some cases are regulated by P availability (Figure S5a-d). *OsPSTOL1* is strongly expressed in the crown region (the base of the shoot where crown roots emerge) (Figure S5e), consistent with the crown root primordia expression of *pOsPSTOL1::uidA* in rice (Gamuyao *et al*. 2012). Although GUS staining was undetectable in other root types or shoot tissues, *OsPSTOL1* mRNA is present in shoots, developing leaves, panicles, roots, and root tips of varieties with a K-allele. This discrepancy between *in planta* GUS staining and mRNA accumulation data may reflect the region used to drive the *uidA* gene, which based on the revised annotation consisted of 272 bp of the promoter followed by a first exon containing a large intron.

*OsPSTOL1-like* genes are associated with development in other species. Sorghum genes(*SORBI_3003G080700, SORBI_3003G263300, SORBI_3003G263400, SORBI_007G032900*) have SNPs associated with root architecture and P uptake (Hufnagel *et al*. 2014; Bernardino *et al*. 2019). These deduced proteins have N-terminal microbial interaction, transmembrane and kinase domains (Figure 4a,b), are expressed in roots and leaves, and induced by pathogens or PAMPs (Figure S5d). The wheat gene (*TaPSTOL1*) is similar protein to *OsPSTOL1* in sequence and domain composition (Milner *et al*. 2018). Its overexpression and knockdown by RNAi influences root and shoot growth and flowering time (Milner *et al*. 2018). Given that *TaPSTOL1* are below the threshold of detection in the Wheat Transcriptome Atlas (Ramírez-González *et al*. 2018) (Figure S5b), its transcript accumulation or protein activity may be limited to specific cells, conditions or accessions.

### 3.4 Ectopic expression of *OsPSTOL1* in bread wheat benefits performance traits

As *OsPSTOL1* has an agronomic benefit in rice and related proteins of sorghum and wheat influence an overlapping set of traits, we hypothesized that *OsPSTOL1* may provide an advantage under low P growth conditions when expressed in wheat. To test this, wheat (cv. Fielder) lines were established that express the *OsPSTOL1* cDNA under the control of the near-constitutive maize *Ubiquitin1* (*Ubi*) promoter. Following initial screening of transgenics, the growth and performance of two independent events (*pUbi::OsPSTOL*1.1 and 1.2) was evaluated relative to a null segregant (Null) in the greenhouse and in a rainfed field trial under replete P conditions (Figure 5). In the greenhouse, we observe a significant benefit in growth and reproduction phenotypes of the transgenics, including greater shoot biomass (*p*<0.04), grain yield (*p*<0.011), and earlier flowering (*p*<1.04×10^−7^) (Figure 5b; Figure S7a, b, and f). *OsPSTOL1* wheat displays more rapid early growth, reaching maturity before the null segregant (Figure 5a). Other traits including 1000 grain weight, grain protein content, total grain nitrogen, heads, tillers, height, and leaf chlorophyll content, were increase in only a single transgenic or unaffected (Figure 5; Figure S7a-l). These results demonstrate ectopic *OsPSTOL1* provides an overall benefit to production without an impact on seed quality traits.

**Figure 5.**
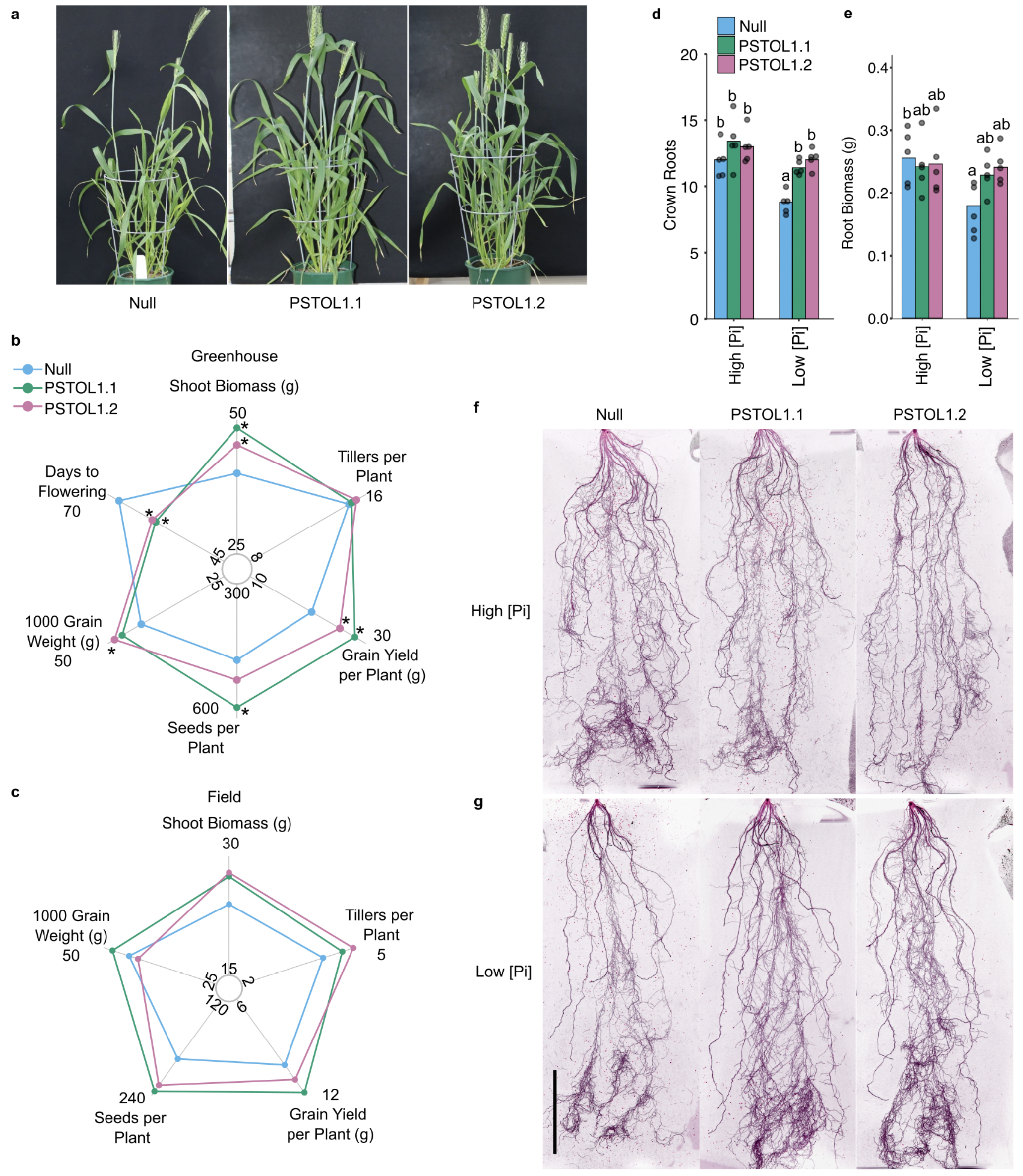
*OsPSTOL1* in wheat can enhance growth in greenhouse and field. **a**, Representative images of three independent *pUbi::OsPSTOL1* transgenic lines and a null segregant at Zadoks stage 65. **b,c**, Yield traits of *pUbi::OsPSTOL1* transgenic lines and their null segregant in a greenhouse (**b**) and a field (**c**) under nutrient replete conditions. For the field, plants were divided into two groups and the average biomass of each group was recorded. The biomass reported is the average of the two plots. Greenhouse: Null, n=10; PSTOL1.1, n=7; PSTOL1.2, n=8; PSTOL1.4, n=14. Field: Null, n=24; PSTOL1.1, n=22; PSTOL1.2, n=21; PSTOL1.4, n=21. **d**, Crown root number and **e**, root biomass of two independent *pUbi::OsPSTOL1* transgenic lines and their null segregant at 28 DAS cultivated under high and low [Pi] conditions in 40 cm tall pots (*n* = 5) in the greenhouse. **f,g** Representative images of the roots of two independent *pUbi::OsPSTOL1* transgenic lines and their null segregant at 28 DAS grown in high (**f**) and low (**g**) [Pi] conditions in 40 cm tall pots. Color has been inverted for visualization. Scale bar, 10 cm. Means significantly different between genotypes are indicated by different letters (*P*< 0.05, analysis of variance (ANOVA) with Tukey’s honest significant difference (HSD) test).

Next, the lines were evaluated in a rain-fed wheat field trial under standard cultivation (P replete) to discern any variation in performance. The results demonstrate a trend of increased shoot biomass (11-16%) and seed number (6-9%) for both transgenics in each replicated plot without an effect on 1000 grain weight (Figure 5c; Figure S8a-f). The 13% increase in grain yield of PSTOL1.1, although not different from the null at P<0.05, exceeds the 1% yield gain per annum typically gained by breeding (Dixon, Braun, Kosina & Crouch 2009; Tester & Langridge 2010; Reynolds *et al*. 2021; Rahman, Crain, Haghighattalab, Singh & Poland 2021). Grain nutrient content was not significantly affected (Figure S9). Above average rainfall during vegetative growth followed by below average rainfall just prior to and at the heading (Figure S10) or variability in the soil microenvironment may have contributed to the trait variability between individual plants and lack of statistical significance. These data indicate that *OsPSTOL1* improves agronomic traits in wheat, but transgenic event by environment interactions influence the overall benefit.

### 3.5 *OsPSTOL1* effects are influenced by P nutrition and developmental stage

Given the performance of PSTOL1.1 and PSTOL1.2 lines under replete P conditions, we sought to determine their performance under P deficiency under controlled greenhouse conditions. We chose to manipulate P availability using a buffered aluminum oxide and sand mixture, shown to more accurately simulate low P stress than unbuffered hydroponics and provide greater reproducibility than comparisons with P replete and deficient soil (Vejchasarn, Lynch & Brown 2016; Hanlon *et al*. 2018). Both lines develop more crown roots and maintain greater root biomass compared to the null segregant under low Pi nutrition (LP, 25 μM), but are indistinguishable when grown under replete P (HP, 436 μM) (Figure 5d-g). P depletion enhances fine root development in all three genotypes at 36 days after sowing (DAS), but this was visibly more pronounced in the transgenics (Figure 6a,b). By 50 DAS under LP, shoot fresh biomass trends higher in PSTOL1.1 and is significantly higher in PSTOL1.2 (Figure 6d). Neither P nor N content are statistically distinguishable between genotypes, although both transgenics trend higher in P (30%) and N (8% [PSTOL1.1] and 29% [PSTOL1.2]) (Figure 6d-e). P content and concentration are lower under LP and decline with age, presumably due to incorporation into shoot tissues, but are indistinguishable between genotypes (Figure S11a-d). The slight difference in phenotype of the two transgenic lines may reflect differences in the level of the *OsPSTOL1* mRNA observed in both root and crown tissue (Figure S12) or spatial-temporal variation in gene activity determined by the transgene insertion site.

**Figure 6.**
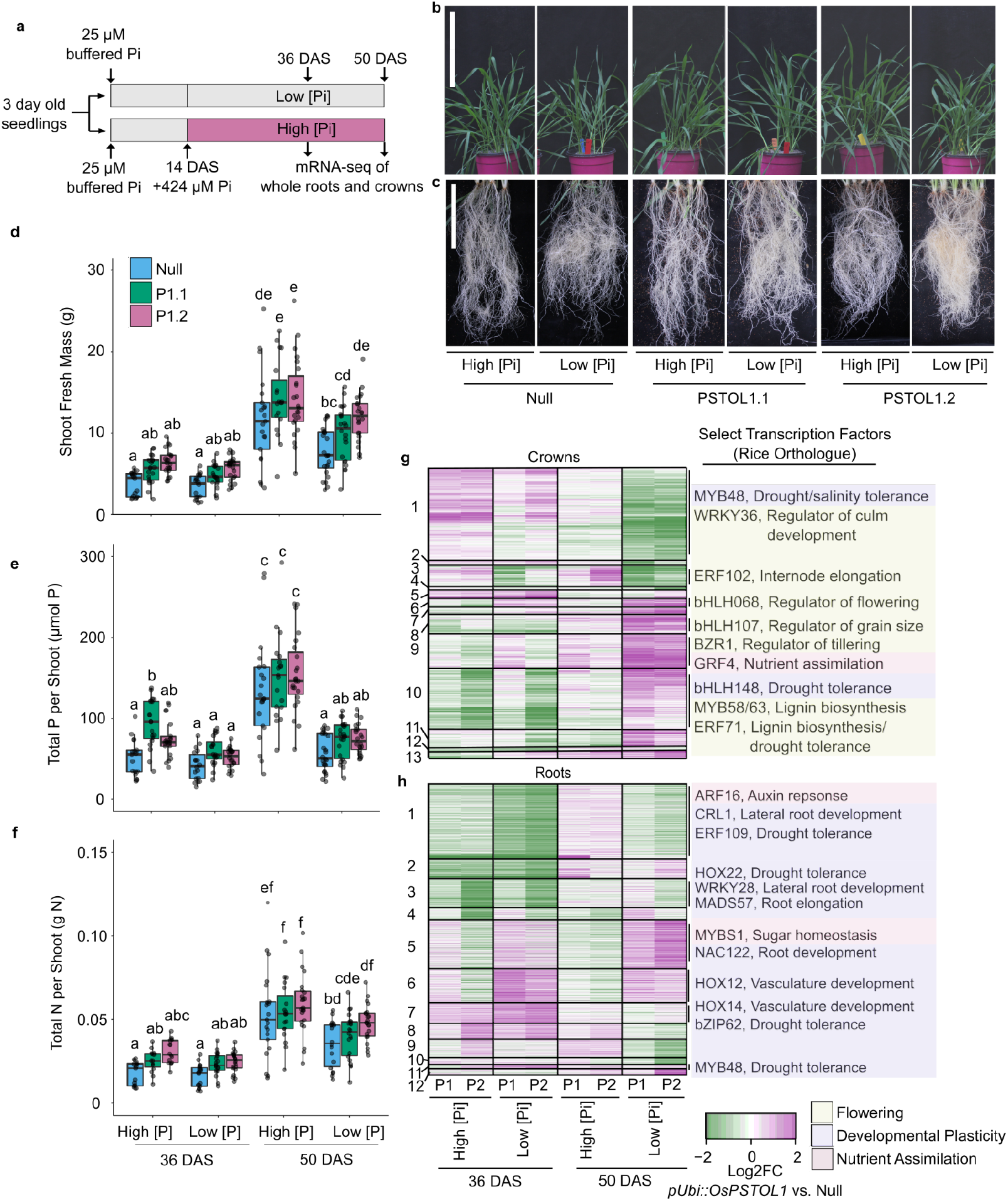
*OsPSTOL1* modestly modifies developmental and P nutrition effects on shoot phenotypes and transcriptional regulator mRNA levels in the shoot crown and roots. **a**, Schematic of the experiment. Three day-old seedlings germinated on plates were transferred to sand culture with 25 μM buffered Pi. Fourteen days after sowing (DAS) high [Pi] (HP) pots were watered every two days with fertilizer containing 424 μM Pi, whereas low [Pi] LP pots were watered with fertilizer containing 0 μM Pi. Root systems and crown regions were harvested at 36 and 50 DAS for mRNA-seq. **b,c** Representative photos of shoots and roots at 36 DAS. **d-f** Shoot fresh mass (**d**), total shoot P content (**e**), and total shoot N content (**f**) from the mRNA-seq experiment tissue; *n*=20. Means significantly different between genotypes are indicated by different letters (*P*< 0.05, analysis of variance (ANOVA) with Tukey’s honest significant difference (HSD) test) Boxplot boundaries represent the first and third quartiles; a horizontal line divides the interquartile range, median. **g,h**, Heatmaps showing differentially regulated genes (DRGs; *P*<0.05, log_2_FC| ≥1) of at least one condition, DAS, tissue or genotype between *pUbi::OsPSTOL1* transgenic line and the null segregant in crowns (**e**) and roots (**f**). P1, PSTOL1.1; P2, PSTOL1.2. Clusters were generated with the Clust package. Selected transcription factors within individual clusters are identified by the rice ortholog name at right.

The root and crown transcriptomes of the three lines were profiled by RNA-seq for the two conditions at 36 and 50 DAS in biological triplicate. Multidimensional scaling plots demonstrate separation by developmental stage, P availability and genotype (Figure S14a,b). Even at this whole transcriptome scale the root transcriptome of the transgenics and Null separate under LP and HP at 36 DAS (panel a, Dimension 2) and in crowns under LP at 50 DAS (panel b, Dimension 2). A total of 1751 elevated and 1658 reduced differentially regulated genes (DRGs) are shared by the two independent transgenic events (*P*<0.05, |log_2_FC| ≥1) across the tissues, stage and treatments (Table S3a). Therefore, we sought genes that were consistently distinguishable in the transgenics relative to null segregant, identifying a greater number of DRGs under LP than HP, with the maximum number in the crown transcriptomes after prolonged cultivation under LP (50 DAS) (Table S3b). This is consistent with greater distinctions in phenotypic traits at 50 DAS. A clustering analysis of the DRGs exposes the degree of uniformity in *OsPSTOL1* effect in the independent transgenic events, as well as distinctions between condition and stage (Figure 6f,g).

To consider common DRGs in both wheat and rice OsPSTOL1 transgenics, we compared the wheat data to the microarray dataset generated from *35S::OsPSTOL1* IR64 rice and its null segregant under low P nutrition (Gamuyao *et al*. 2012). Of the 23 DRGs identified in rice, 11 correspond to wheat orthologs of DRGs in OsPSTOL1 wheat under at least one condition (Figure S14, Table S3c). Orthologs of the transcription factor *DELAY OF ONSET OF SENESCENCE* (*OsDOS*) that limits senescence, S-adenosyl methionine decarboxylase (*SAMDC2*) that is involved in polyamine biosynthesis, and several others stand out as similarly regulated in OsPSTOL1 transgenics of both species, indicating similar downstream effects.

### 3.6 Transcription factors related to root development, drought stress, and flowering are differentially regulated in roots and crowns of OsPSTOL1 wheat

Transcription factors were subset from the DRG clusters to gain insight into developmental or environmental response regulation conferred by *OsPSTOL1*. Due to limited analyses of transcription factors in wheat, we focused our analysis on known function of the orthologous genes in rice. A number of the DRG are transcription factor associated with root development and environmental responses (Figure 6f,g; Figure S15a,b; Table S3d,e). These include orthologs of *OsHOX12/14*, involved in organ development (Shao *et al*. 2018) and *OsNAC122*/*10* influencing root diameter and drought tolerance (Jeong *et al*. 2010). Homologs of *OsCRL1*, involved in lateral root and crown initiation (Inukai *et al*. 2005), are down-regulated in OsPSTOL1 roots at 36 DAS but up-regulated in the crown at 50 DAS. These results indicate that the transgenics differentially regulate mRNAs associated with root initiation. Other DRGs encode Sugars Will Eventually Be Exported Transporters (SWEET), Trehalose-6-phosphate synthases (TPS) and phosphatases (TPP) that regulate the allocation of carbohydrates from photosynthetic source organs to sink tissues, such as roots, crowns or developing seed (Paul, Gonzalez-Uriarte, Griffiths & Hassani-Pak 2018) (Figure S16; Table S3f). *SWEET* mRNAs are higher in the transgenic roots HP particularly at 50 DAS, suggesting there may be greater resource allocation to continue root development. OsPSTOL1 may directly or indirectly influence the partitioning of carbohydrates from source leaves to roots.

OsPSTOL1 transgenics showed small but beneficial alterations in a number of developmental traits including a reduction in days to flowering coupled with greater shoot biomass and grain yield (Figure S7a,b,f). Indeed, OsPSTOL1 significant influences transcript levels of orthologs of a number of developmental regulators in both transgenics (Figure 6f,g; Figure S15). These include wheat orthologs of *OsbHLH068, OsBZR1, OsGRF4*, Os*HOX12* and *OsbHLH107*, associated with internode elongation, tillering, nutrient assimilation and yield (Li *et al*. 2018), flowering, and panicle development in rice (Tong *et al*. 2009; Gao, Fang, Xu, Wang & Chu 2016; Chen *et al*. 2017; Shao *et al*. 2018; Yang *et al*. 2018). The differential regulation of genes involved in reproduction are of interest as P deficiency typically delays flowering in rice and wheat (Rodríguez, Pomar & Goudriaan 1998; Ye *et al*. 2019). The DRGs include several orthologs of genes implicated in ABA signaling and drought resilience, most of which show strong upregulation at 50 DAS in crowns. The elevation of these regulators could reflect the promotion or an advancement in development by ectopic OsPSTOL1 expression. Of particular note is the downregulation of orthologs of *OsARF16* and *OsWRKY28*, involved in changes in root architecture under P deficiency (Shen *et al*. 2013; Wang *et al*. 2018a). This led us to consider that P sensing or mobilization may be altered in OsPSTOL1 transgenics.

### 3.7 Evolutionarily conserved core low P response pathway is induced earlier in P-starved OsPSTOL1 wheat

The prior DNA microarray analysis of *35S*::*OsPSTOL1* and *OsPSTOL1* near isogenic lines of rice provided no indication of an effect on P sensing or response after prolonged P deficiency (Pariasca-Tanaka, Satoh, Rose, Mauleon & Wissuwa 2009; Gamuyao *et al*. 2012). Taking advantage of the inventory of low P responsive genes of rice (Oono *et al*. 2013a), we observe progressive elevation of the orthologous low marker genes in roots at 36 DAS, followed by greater induction at 50 DAS in roots and crowns (Figure S17a,b; Table S3g,h). A subset of these mRNAs rises early in transgenic roots at 36 DAS (clusters 4, 8 and 9, Figure 7a, S17b). The precocious elevation of these transcripts led us to systematically analyze genes in the low P signaling and response pathway (Figure 7). Our survey starts with the PHR MYB-type transcription factors that control downstream low P responses. We observe that *PHR3* orthologs are elevated in roots at 36 DAS, consistent with upregulation of downstream low P response genes. However, PHRs are tightly controlled by SPXs, which post-translationally repress their activity (Poirier 2019; Wang *et al*. 2009). Early and pronounced upregulation of *SPX*s suggests that PHR upregulation is counterbalanced in the transgenics. PHR activity also serves to limit levels of P transporters. PHRs upregulated *IPS1*, a noncoding RNA that acts as a molecular sponge of *miR399* to limit accumulation of *PHO2* (*PHOSPHATE 2*) mRNA, encoding an E3 ligase that catalyzes the degradation of the P transporters. The greater reduction in *PHO2* mRNA, despite a significant elevation of *IPS1* particularly in PSTOL1.2, suggests that either the level or spatial distribution of *IPS1* is insufficient to curb *miR399* activity. It also suggests that turnover of P transporter may be reduced in the transgenics.

**Figure 7.**
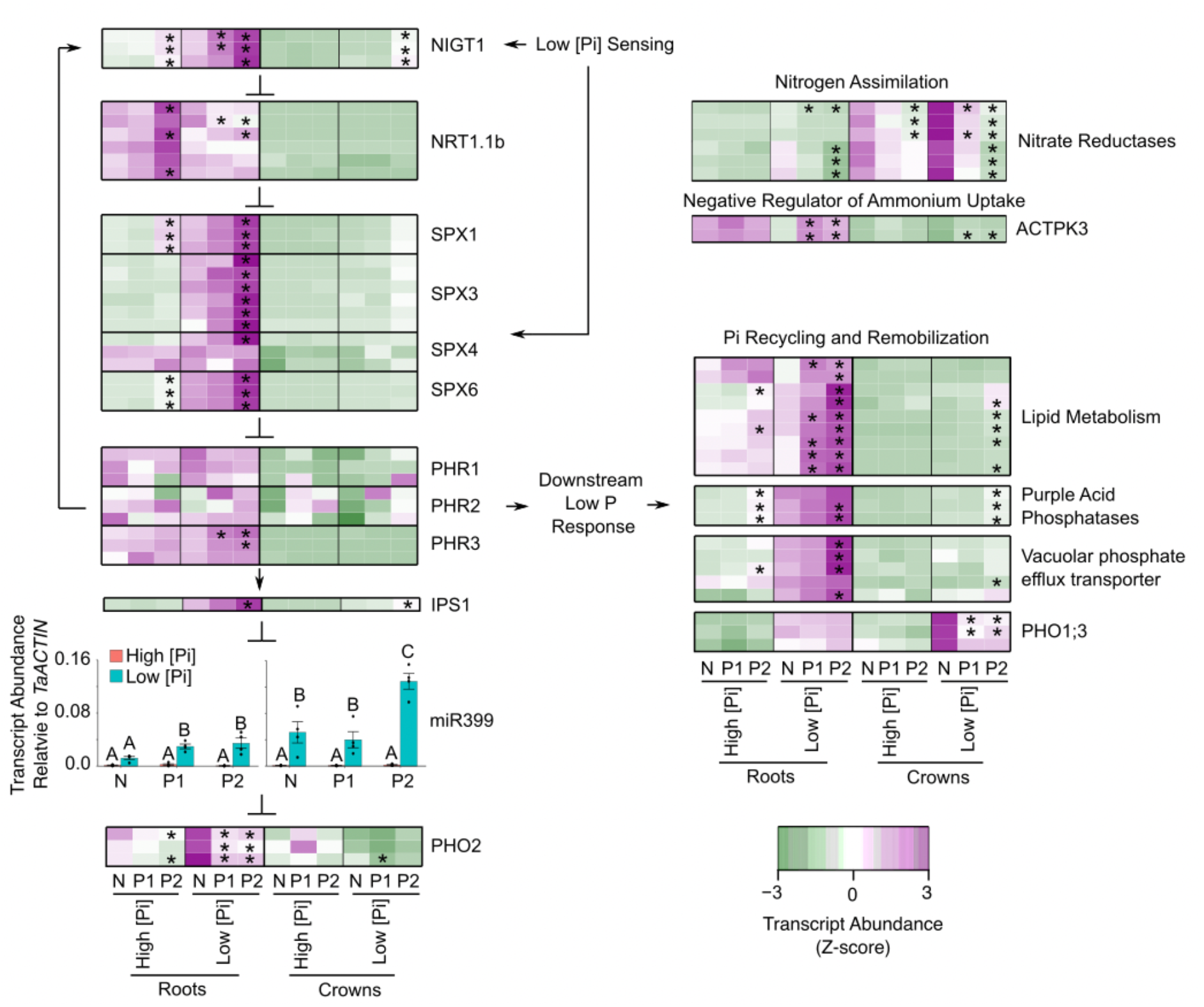
*OsPSTOL1* wheat precociously elevates core low P response pathway and Pi recycling transcripts under P deficiency. Transcript abundance normalized by Z-score of the core low P response pathway and downstream genes at 36 DAS in roots and crowns in *pUBI::OsPSTOL1* transgenics (P1 = *pUBI::OsPSTOL1.1*; P2 = *pUBI::OsPSTOL1.2*) and their null segregant (N). Each row is one gene; homologs are grouped. An asterisk indicates significantly different than the null segregant at the same P status and organ (*P*<0.05). miR399 transcript was measured with qRT-PCR relative to *TaACTIN*.n=4. Means significantly different between genotypes and treatments within roots or crowns are indicated by different letters (*P*< 0.05, analysis of variance (ANOVA) with Tukey’s honest significant difference (HSD) test). Genes shown are differentially regulated in at least one transgenic line in at least one condition in *pUbi::OsPSTOL1* transgenic lines compared to null segregant (*P*<0.05), with differentially regulated genes indicated with an asterisk.

These data indicate both elevation and dampening of the typical P-starvation response in the transgenics. On one hand, PHR-regulated genes are associated with P recycling and remobilization are significantly elevated at 36 DAS in both lines (Figure 7; Table S2h). These include enzymes of sulfolipid and galactolipid metabolism reported to be low P-induced in wheat (Oono *et al*., 2013b; Walkowiak *et al*., 2020). Remarkably, there is precocious activation of a low P response signature in roots growing under P replete conditions at 36 DAS in the PSTOL1.2 line. This includes significant precocious elevation of ten P transporter mRNAs (Figure S17c). On the other hand, despite the changes in *PHR* mRNAs and the *miRNA399*-*IPS1*-*PHO2* module components at 36 DAS, mRNAs encoding many known P transporter orthologs are not differentially regulated under LP in the wheat transgenics as observed in *OsPSTOL1* rice (Gamuyao *et al*. 2012). Finally, *PHO1;3* mRNA, encoding the P starvation-induced transporter involved in root to shoot partitioning of P is significantly lower in the transgenic crowns.

The activation of the low P pathway is reciprocally joined by the inhibition of N uptake (Hu *et al*. 2019). Consistent with possible heightened perception of P depletion, mRNAs encoding orthologs of the N response sensor and transporter (transceptor) *NRT1.1b* and N assimilation enzymes (nitrate reductases) are lower in roots and crowns of both transgenics under P starvation (Figure 7). *NIGT1s*, encoding a negative regulator of *NRT1.1b*, are upregulated in OsPSTOL1.2 under HP and LP at 36 DAS. This is consistent with elevated *SPX* but not that of *PHR* and *IPS1* mRNAs. It is possible that SPX levels are limited by a NRTL.1B-interacting protein 1 E3 ligase, as in P-deprived rice (Hu *et al*. 2019). These results support the conclusion that *OsPSTOL1* accentuates low P sensing even under replete P, predisposing roots to acquire and remobilize P resources proactively but with pronounced counterbalancing, that might be less complex if analyzed in roots of specific type or individual cells.

## 4 DISCUSSION

*OsPSTOL1* is variably present in both domesticated rice and its wild relatives due to multiple independent gene mutation or deletion events (Pariasca-Tanaka *et al*. 2014; Vigueira *et al*. 2016), demonstrating it is non-essential but could be beneficial in certain environments. Low frequency of a functional *OsPSTOL1* in east Asia is at least partially due to a gene deletion that occurred after the tropical-japonica split. It is not clear if this loss reflects selection or the genetic bottleneck that the founders of the temperate population passed through (Gutaker *et al*. 2020). Our biogeographical survey of landraces and accessions confirm *OsPSTOL1* is depleted in *japonica* and *indica* landraces adapted to irrigated lowlands but is enriched in *japonica* varieties cultivated in rainfed uplands prone to low nutrient availability and drought (Haefele *et al*. 2014). To date, studies showing a benefit of *OsPSTOL1* with low P input were performed under aerobic (non-waterlogged) conditions, with an increased benefit under simultaneous water deficiency (Wissuwa & Ae 2001a b; Pariasca-Tanaka *et al*. 2009; Chin *et al*. 2010; Gamuyao *et al*. 2012), common in rainfed ecosystems after seedling establishment (Haefele *et al*. 2014). We find that rice varieties of west Africa and Madagascar have high frequencies of *OsPSTOL1*, where cultivation is almost exclusively rainfed and soil nutrient quality is often low (Haefele *et al*. 2014). Thus, although a cost of the presence of OsPSTOL1 in paddy cultivation has not been tested, this gene is likely to provide an adaptive benefit in nutrient poor and dry aerobic soils.

Our phylogenetic analyses indicate *OsPSTOL1* is truncated to a solo kinase domain relative to related genes found across Gramineae crops, which typically possess a N-terminal region associated with molecular interactions with microbes, followed by transmembrane and kinase domains. Microbes or PAMPs rather than P availability induce many of these genes. Our transcript revaluation finds that *OsPSTOL1* encodes the conserved transmembrane sequence within its 5’ UTR, confirming its origin by truncation but also raising the possibility that use of a non-AUG start codon could result in synthesis of a kinase with a N-terminal transmembrane domain. Indeed, genetic variation in four sorghum Os*PSTOL1-*like genes correlate with root architecture and low P tolerance (Hufnagel *et al*. 2014; Bernardino *et al*. 2019); unlike *OsPSTOL1* and *TaPSTOL1*, these genes have N-terminal and transmembrane domains and display transcript induction by pathogens. This emphasizes that this family of modular genes may be valuable subjects for improving root architecture to enhance nutrient acquisition, including P and N, in multiple Gramineae, motivating our evaluation of *OsPSTOL1* in bread wheat.

Through heterologous expression in wheat, we show that *OsPSTOL1* acts on evolutionarily conserved pathways that influence developmental and environmental responses. Modest benefits to seed number and yield, were evident in *OsPSTOL1* wheat cultivated with fertilization in the greenhouse and rainfed field, without a reduction in 1000 grain weight, which can occur when seed number increases due to finite C and N resources delivered to seed. This indicates greater efficiency in nutrient acquisition or mobilization. These phenotypes are accompanied by elevated levels of transcription factors associated with growth, flowering, and yield in the crown tissue. Under P deficiency, root and shoot growth is enhanced in the transgenics relative to the null segregant, along with a trend towards greater P and N content in shoot tissue under both deficient and replete P conditions that increases towards bolting. Notably, cumulative benefits on P uptake in OsPSTOL1 rice have only been observed late in development, typically after 70 days (Wissuwa & Ae 2001b; Pariasca-Tanaka *et al*. 2009).

To better understand the molecular function of OsPSTOL1 in wheat, we monitored root and crown transcriptomes during development and in response to P nutrition. We find that *OsPSTOL1* precociously amplifies the core low P sensing and response of roots, enhancing early elevation of transcripts associated with P remobilization and recycling. After prolonged P deficiency, OsPSTOL1 roots and crowns have modified levels of mRNAs encoding transcription factor that control nutrient assimilation, sugar homeostasis and reproductive development. The transcriptome data suggest OsPSTOL1 may limit N uptake at 36 DAS under low Pi, however N accumulation consistently tracks with biomass accumulation in the transgenics, possibly due the production of fine roots that increase total root surface area. Higher resolution analyses, such as single cell RNA-sequencing and spatial transcriptomics will be necessary to decipher the complex role of OsPSTOL1 in low P sensing and response indicated by the organ-level transcriptome study.

The *OsPSTOL1* effect includes elevated expression of mRNAs encoding transcription factors involved in root development and drought tolerance. Among these genes are several *HOX*s, also differentially regulated in *OsPSTOL1* rice transgenics (Gamuyao *et al*. 2012), suggesting these developmental regulators are conserved downstream targets of OsPSTOL1 and may determine the plasticity associated with vigor in dry and low nutrient ecosystems. This and the precise cellular function of OsPSTOL1 deserve further investigation, as contrasting root system architecture is generally associated with robust P acquisition by shallow roots with dense hairs (Lynch 2011) and water acquisition under deficit by thin deep lateral roots (Klein, Schneider, Perkins, Brown & Lynch 2020). It seems likely that the selection of *OsPSTOL1* in low nutrient rainfed ecosystems with recurrent dry periods reflects appropriate regulation of developmental plasticity and P sensing that favor nutrient and moisture capture.

## 5 CONCLUSIONS

This study provides new insights into the roles of *OsPSTOL1* as a valuable adaptive gene by linking information on its biogeographical distribution across rice subspecies and ecosystems, clarifying protein evolution, and structural features. It also demonstrates cumulative benefits of *OsPSTOL1* ectopic expression in wheat on root systems, yield traits, and adaptive gene regulation. To further understand the agronomic costs and benefits of *OsPSTOL1*, it will be valuable to address potential interactions in terms of drought, mycorrhizal fungi, and the acid soil syndrome that commonly co-occur with P deficiency.

## ACKNOWLEDGEMENTS

We thank Alexander Borowsky and Garo Akmakjian of the Bailey-Serres group, Melissa Pickering, Laura Short and Christine Tritterman of the Roy group, Eduardo Gaxiola, Ravi Valluru, and Louis Santiago for thoughtful discussion. This research was supported by an International Wheat Yield Project grant to S.R., J.B.-S. and S.H., US National Science Foundation grants (IOS-1238243, IOS-1856749, IOS-1936492) and a UC MacArthur Foundation Chair award to J.B.-S; Grains Research and Development Council (ACP0009) to S.R.; and Rothamsted Research strategic funding from the Biotechnological and Biological Sciences Research Council of the United Kingdom and the Designing Future Wheat (DFW) Strategic Programme (BB/P016855/1) to S.H..

## AUTHOR CONTRIBUTIONS

A.K., M.A.L., K.Y., S.R. and J.B.-S. conceived and designed experiments; A.K., M.A.L., K.Y., T.R., and S.B, M.J.P. and S.R performed experiments; A.K., M.A.L., K.Y., T.R., S.R., S.H. and J.B.-S. analyzed data; A.K. and J.B.-S. wrote the manuscript; A.K., M.A.L., M.J.P., S.H. and S.R. edited the manuscript.

## CONFLICT OF INTEREST

The authors declare no conflict of interest.

## Supporting Information: Tables

Table S1 Publicly available mRNA-seq datasets, primers, and fertilizers used in this study

Table S2 *Oryza* genomic resources and analyses

Table S3 Transcriptomic analysis of *pUbi::OsPSTOL1* transgenics of wheat

## Supporting Information: Methods (see end of document)

## Supporting Materials

## Figures and Methods

**Figure S1.**
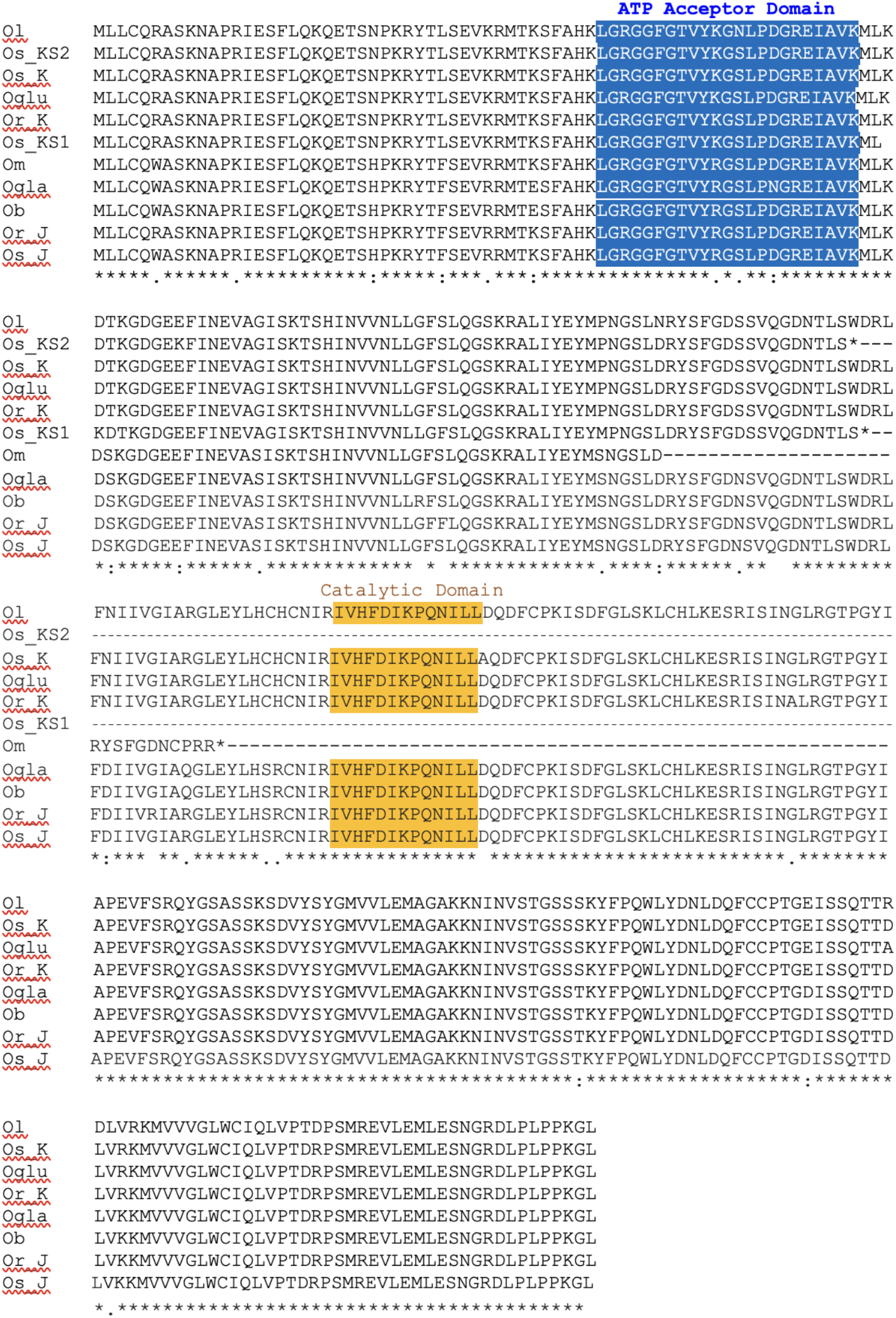
Amino acid sequence alignment of *OsPSTOL1* with the orthologous gene of AA genome *Oryza*. Highlighted in blue is the ATP acceptor domain and highlighted in orange is the catalytic domain. Oglu, *O. glumaepatula*; Os_K, *O. sativa* K-allele; Os_KS2, *O. sativa* KStop2 allele; Or_K, *O. rufipogon* K-allele; Ol, *O. longistaminata*; Om, *O. meridionalis;* Ogla, *O. glaberrima*; Ob, *O. barthii*; Or_J, *O. rufipogon* J-allele; Os_J, *O. sativa* J-Allele; Os_KS1, *O. sativa* KStop1 Allele. KStop1 and KStop2 result from single nucleotide mutations at 410 and 411 of the coding sequence, respectively that result in an in-frame stop codon shown as an asterisk in the protein sequence. Related to Figure 1.

**Figure S2.**
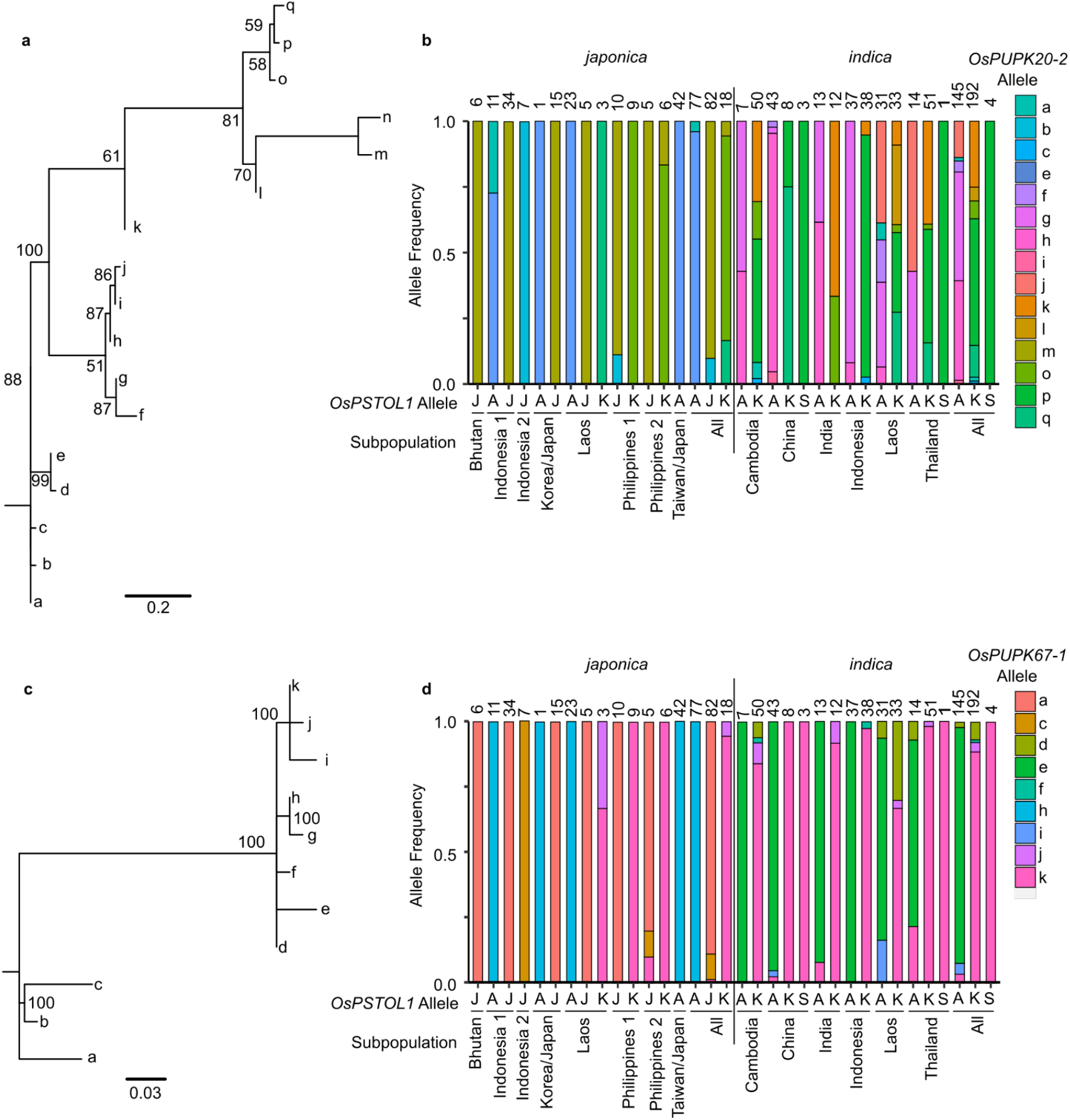
Allelic variation at flanking loci is linked to allelic variation at the *OsPSTOL1* locus. **a**, Haplotypes of select *japonica* subpopulations for the *OsPUPK20-2* and *OsPSTOL1* loci on chromosome 12. The distance between *OsPupk20-2* and *OsPSTOL1* is predicted from the Kasalath *Pup1* locus sequence. Genes are not drawn to scale. **b,c**, Phylogenetic trees of *OsPUPK20-2* (**a**) and *OsPUPK67-1* (**c**) alleles based on SNPs in the coding region. Only alleles present in at least ten varieties are shown. Node labels indicate the percentage of 1000 bootstraps. **d,e**, Distribution of *OsPUPK20-2* (**b**) and *OsPUPK67-1* (**d**) alleles across subpopulations of rice landraces. A, Absent haplotype; J, J-Allele; K, K-allele; S, KStop2. Related to Figure 2 and supported by Table S2f.

**Figure S3.**
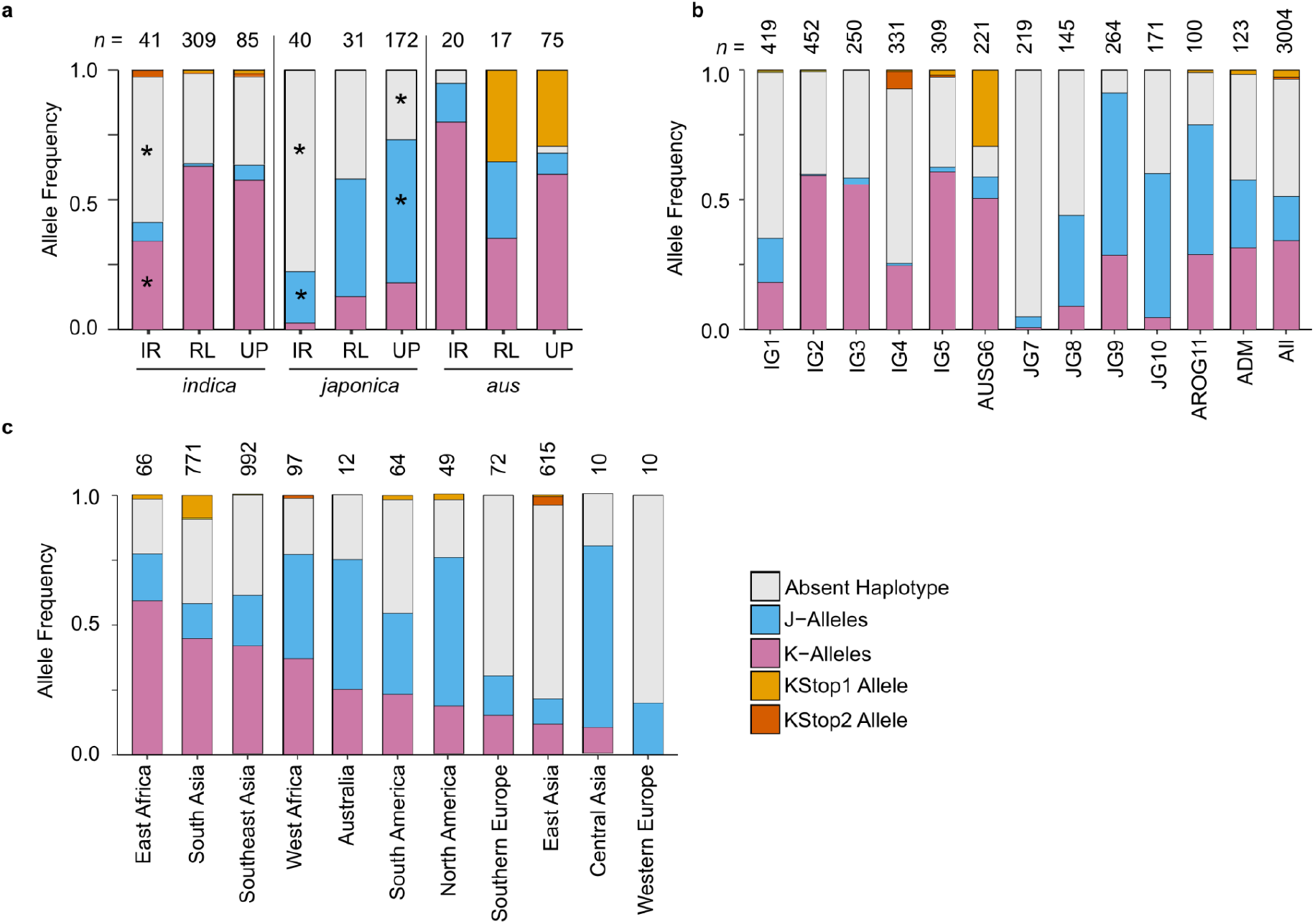
Absence of *OsPSTOL1* is more common in *indica* and *japonica* landraces grown in irrigated lowlands. **a**, Distribution of *OsPSTOL1* alleles in landraces by subspecies and rice growing ecosystems. Asterisk indicates functional *OsPSTOL1* (J- and K-allele) or loss-of-function of *OsPSTOL1* (KStop1, Kstop2, and Absent Haplotype) are statistically different than expected by *Chi*-square test (* indicates *P*<0.05). IR, irrigated lowland; RL, rainfed lowland; UP, rainfed upland. Too few deepwater, swamp water and tidal water landraces were surveyed to make definitive conclusions. **b**, Distribution of *OsPSTOL1* haplotypes based on the SNP groups identified in the 3000 Rice Genome dataset. **c**, Distribution of *OsPSTOL1* haplotypes based on geographic region defined by FAOSTAT. Countries with fewer than 10 varieties sampled were excluded. Varieties with an ambiguous haplotype or multiple haplotypes were excluded. Related to Figure 3.

**Figure S4.**
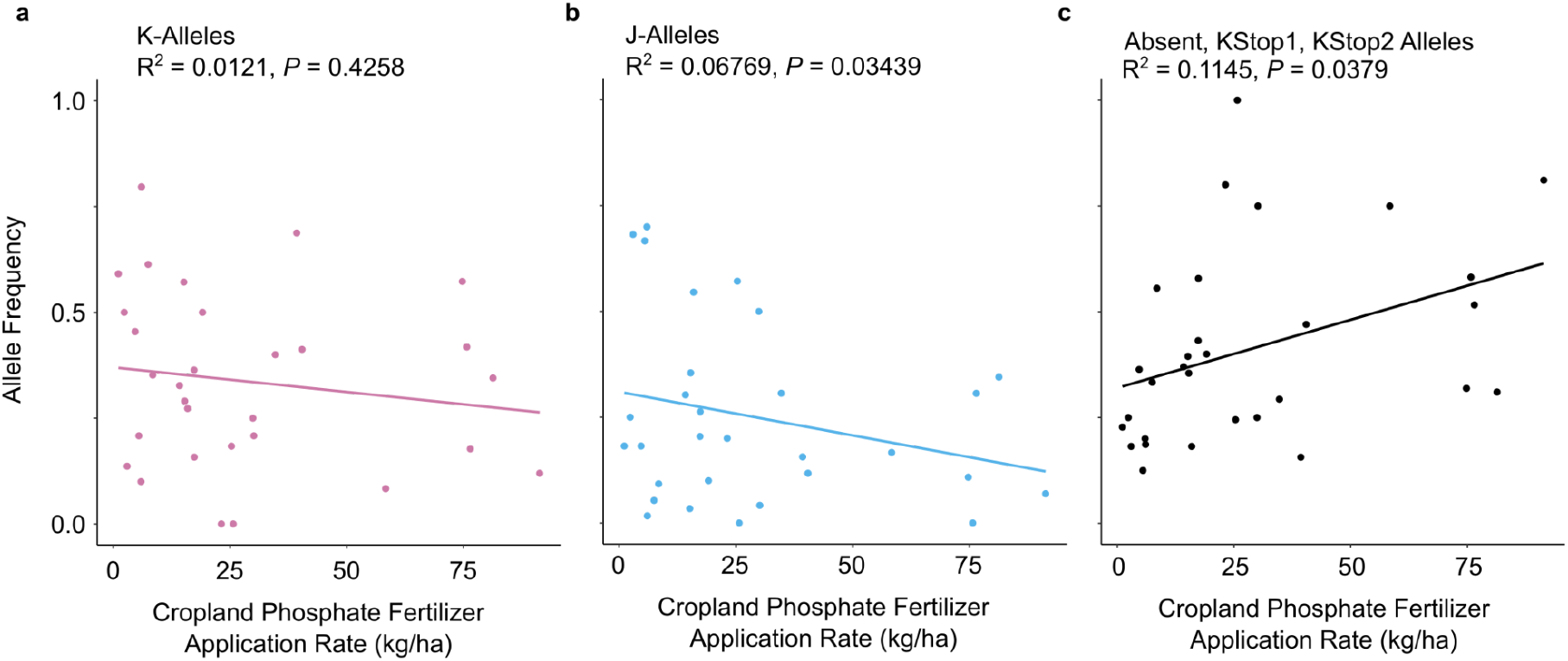
Loss of *OsPSTOL1* weakly correlates with cropland phosphate fertilizer application rate. **a-c**, Correlation of the frequency of K-allele (**a**), J-allele (**b**), and three loss of function alleles (**c**) with the cropland P fertilizer application rate in each country (Roser & Ritchie 2013). R^2^ and best fit line calculated using linear regression. Related to Figure 3.

**Figure S5.**
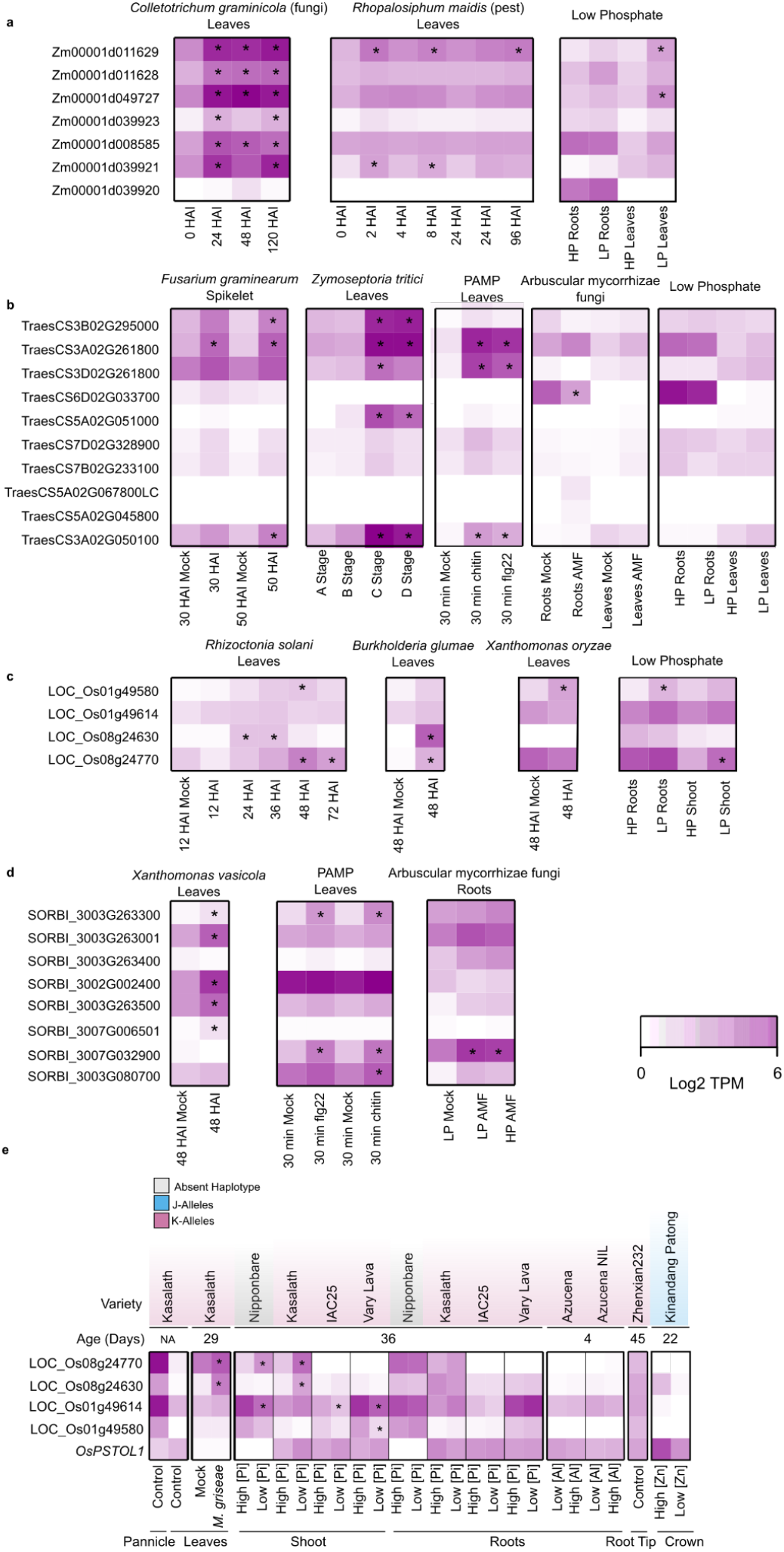
*OsPSTOL1*-like genes are generally induced by pathogens and *OsPSTOL1* is broadly expressed in the roots, shoots, and panicles but shows less conditional regulation than related genes. **a-d**, Transcript abundance of *OsPSTOL1*-like genes in maize (**a**), wheat (**b**), rice (**c**), and sorghum (**d**) under various biotic and low P stresses and in various organs and conditions in rice (**e**). HAI, hours after inoculation; HP, high [Pi].; LP, low [Pi]; AMF, arbuscular mycorrhizal fungi. [Pi], phosphate; *M. griseae, Magnaporthe griseae*; Al, aluminum; Zn, Zinc. See Table S1a for details of the public data sources. Asterisks indicate significantly different from the control condition (*P*<0.05, |log_2_FC| ≥1).

**Figure S6.**
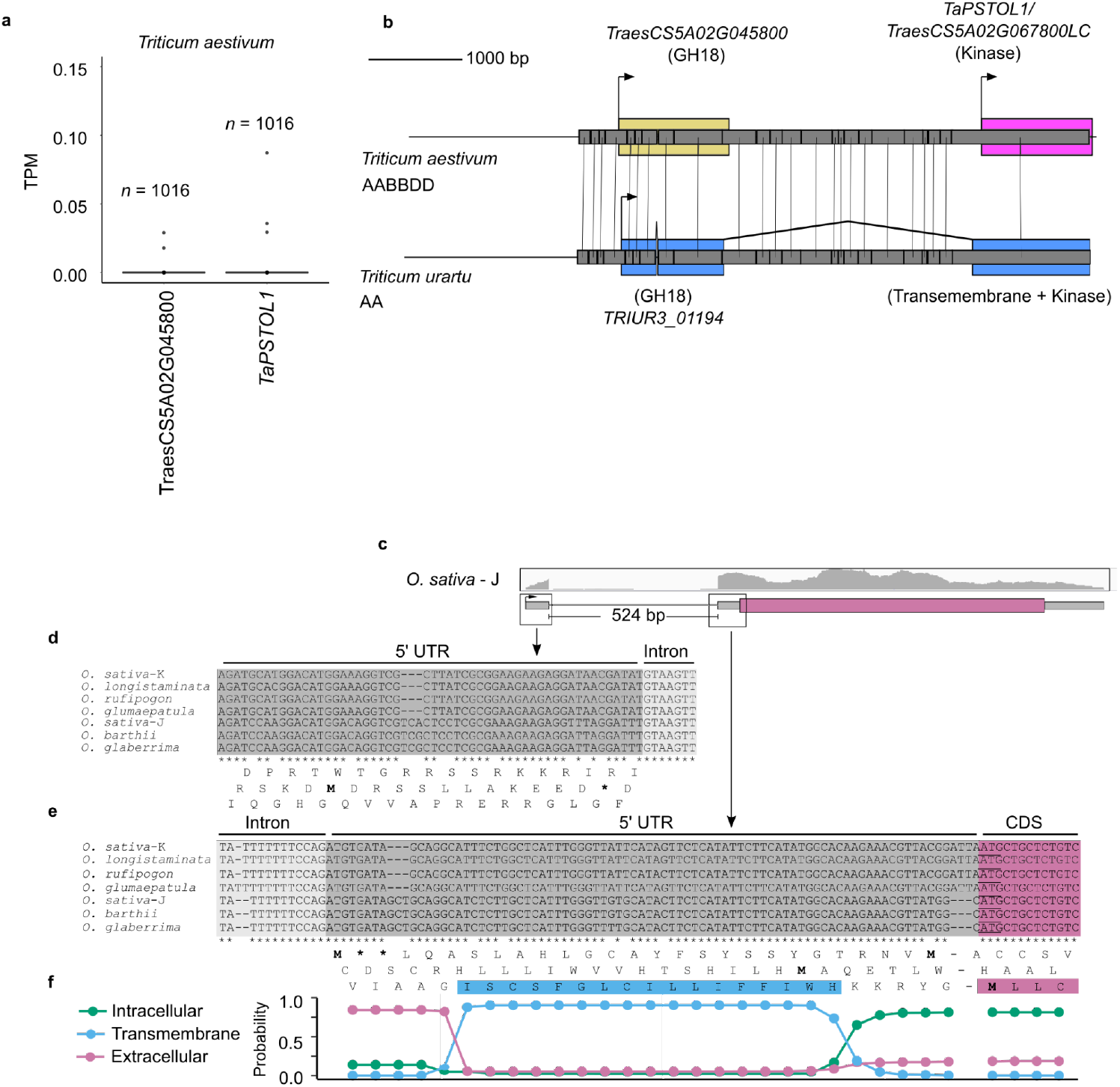
Evidence for limited expression of TaPSTOL1 and origin involving truncation of the kinase-domain only OsPSTOL1 and TaPSTOL1. **a**, *TaPSTOL1* (*TraesCS5A02G067800LC*/*Traes_5AS_AA3DC6A5F*) and the most closely related gene *TraesCS5A02G045899* are detected below the threshold in nearly all samples in the wheat transcriptome atlas(Ramírez-González *et al*. 2018). **b**, Nucleotide sequence alignment of the wheat gene *TaPSTOL1* and its upstream gene *TraesCS5A02G045899* to the genomic region of the *Triticum urartu TRIUR3_01194*. Gray rectangles between the two loci indicate segments of conserved sequence, with variant nucleotides shown as vertical black lines. GH18, glycoside hydrolase 18 domain. **c**, Alignment of RNA-seq reads from publicly datasets to *O. sativa OsPSTOL1* J-allele. mRNA-seq derived from the crown was used for alignment. Magenta predicted coding sequence; gray box, predicted 5’ untranslated region (UTR), and black horizontal line for an intron deduced from RNA-seg data are supported by canonical intron splice donor and acceptor sites. RNA-seq reads were visualized with a genome browser and reads mapped in grey are displayed within a black horizontal box, placed above the gene structure diagram. **d,e**, Nucleotide sequence alignment of exon 1 and the 5’ end of the intron positioned within the 5’ UTR (**d**) and the 3’ end of the intron and 5’ end of exon 2 that precedes the start codon (**e**) of *OsPSTOL1*. Translation of the three reading frames for *O. sativa-J* are shown for comparison to the K-allele data in Figure 4f. Asterisk indicates a stop codon. Blue highlighted region indicates the putative transmembrane domain that resides within the 5’ UTR of the gene. **f**, Transmembrane domain prediction calculated using Phobius for the amino acid sequence in the third frame of (**e**). *OsPSTOL1* mRNA translation is predicted to initiate downstream of this domain. See Table S1a for details of the datasets and accessions surveyed.

**Figure S7.**
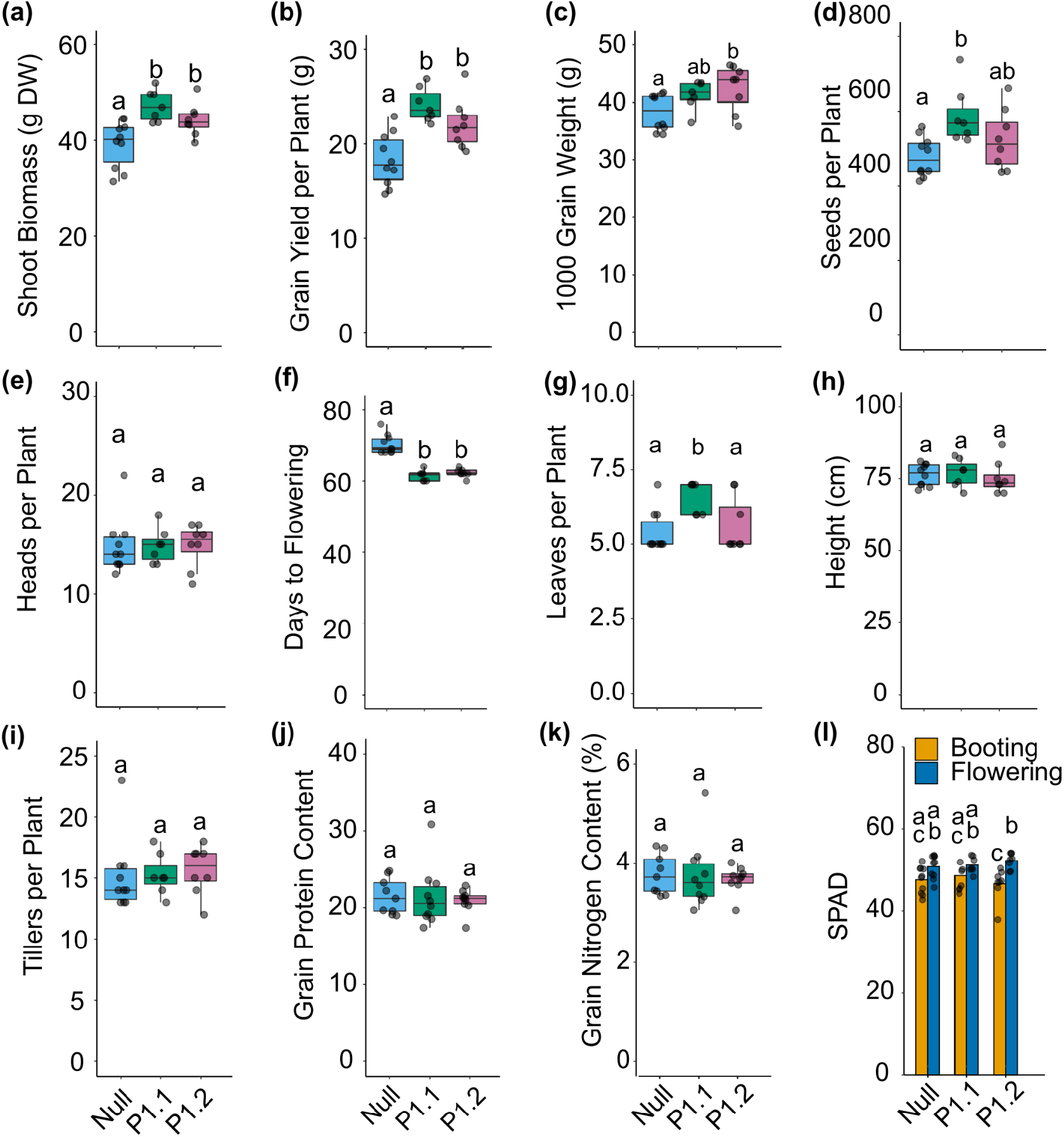
Biomass and yield trait phenotyping of *pUbi::OsPSTOL1* transgenics and their null segregant under P replete greenhouse conditions. **a-k**, Shoot biomass (**a**), grain yield per plant (**b**), 1000 grain weight (**c**), seeds per plant (**d**), heads per plant at maturity (**e**), days to flowering (**f**), leaves per plant (**g**), height at maturity (**h**), tillers per plant at maturity (**i**), grain protein content (**j**), and grain nitrogen content (**k**) of *pUbi::OsPSTOL1* transgenic lines and their null segregants grown in a greenhouse in south Australia under nutrient replete conditions. Boxplot boundaries represent the first and third quartiles; a horizontal line divides the interquartile range, median. **l**, SPAD meter readings of chlorophyll of mature leaves at booting and flowering stage. Means significantly different between genotypes are indicated by different letters (*P*<0.05, analysis of variance (ANOVA) with Tukey’s honest significant difference (HSD) test). Related to Figure 5.

**Figure S8.**
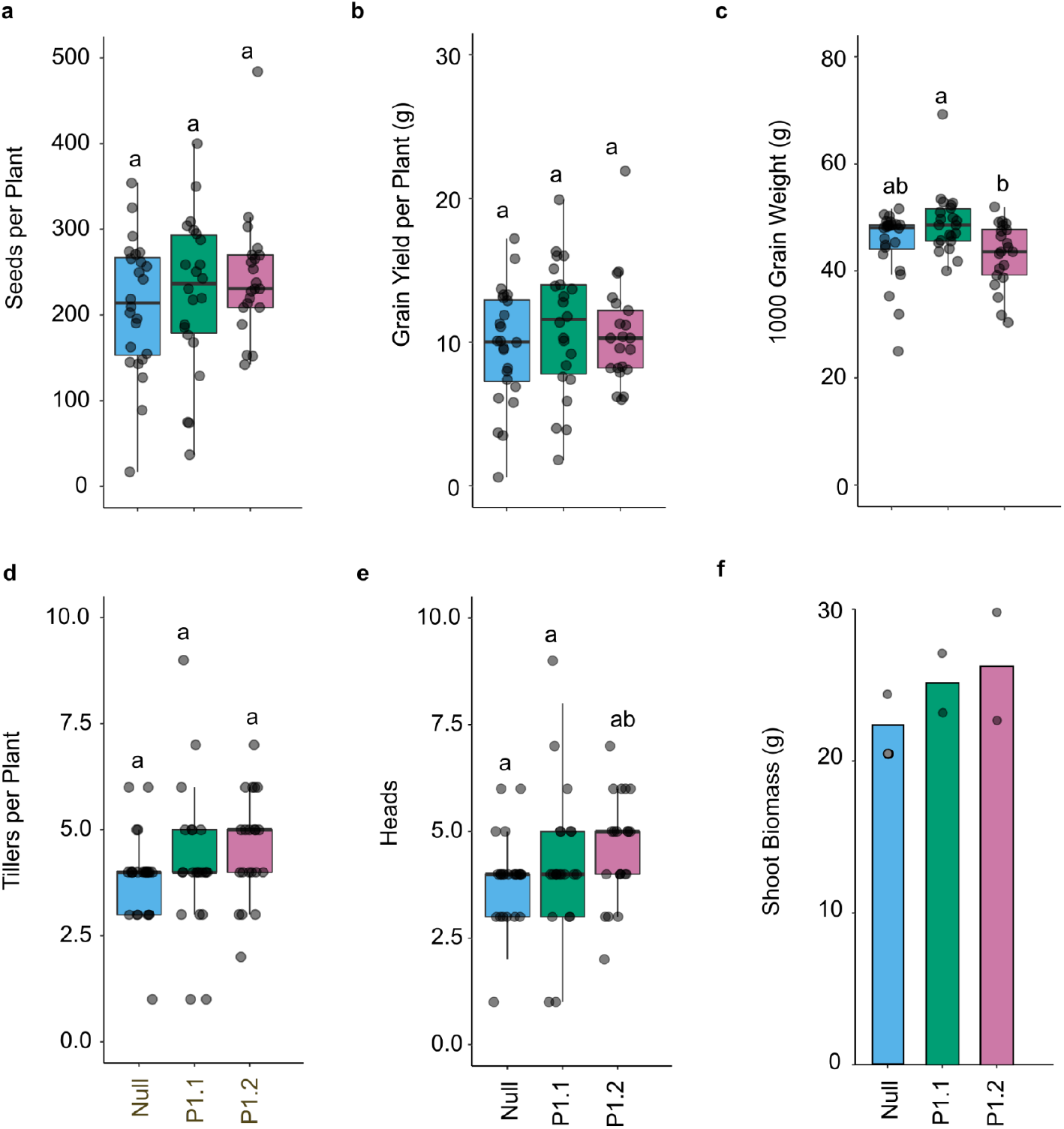
Yield Traits of *pUbi::OsPSTOL1* transgenics and their null segregant grown in the field. Characteristics of the productivity of wheat cultivated in a field trial at Glenthorne farm in South Australia under nutrient replete conditions. **a-f**, Total seeds per plant (**a**), grain yield per plants (**b**), 1000 grain weight (**c**), tillers per plant at maturity (**d**), number of heads at maturity (**e**), and shoot biomass (**f**) of *pUbi::OsPSTOL1* transgenic lines and their null segregants grown in a field in South Australia under nutrient replete conditions. For shoot biomass, plants were divided into two groups and the average biomass of each group was recorded. The biomasses reported are the averages of the two plots. Null, n=24; PSTOL1.1, n=22; PSTOL1.2, n=21; PSTOL1.4, n=21. Means significantly different between genotypes are indicated by different letters (*P*<0.05, ANOVA with Tukey’s honest significant difference (HSD) test). Boxplot boundaries represent the first and third quartiles; a horizontal line divides the interquartile range, median. Related to Figure 5.

**Figure S9.**
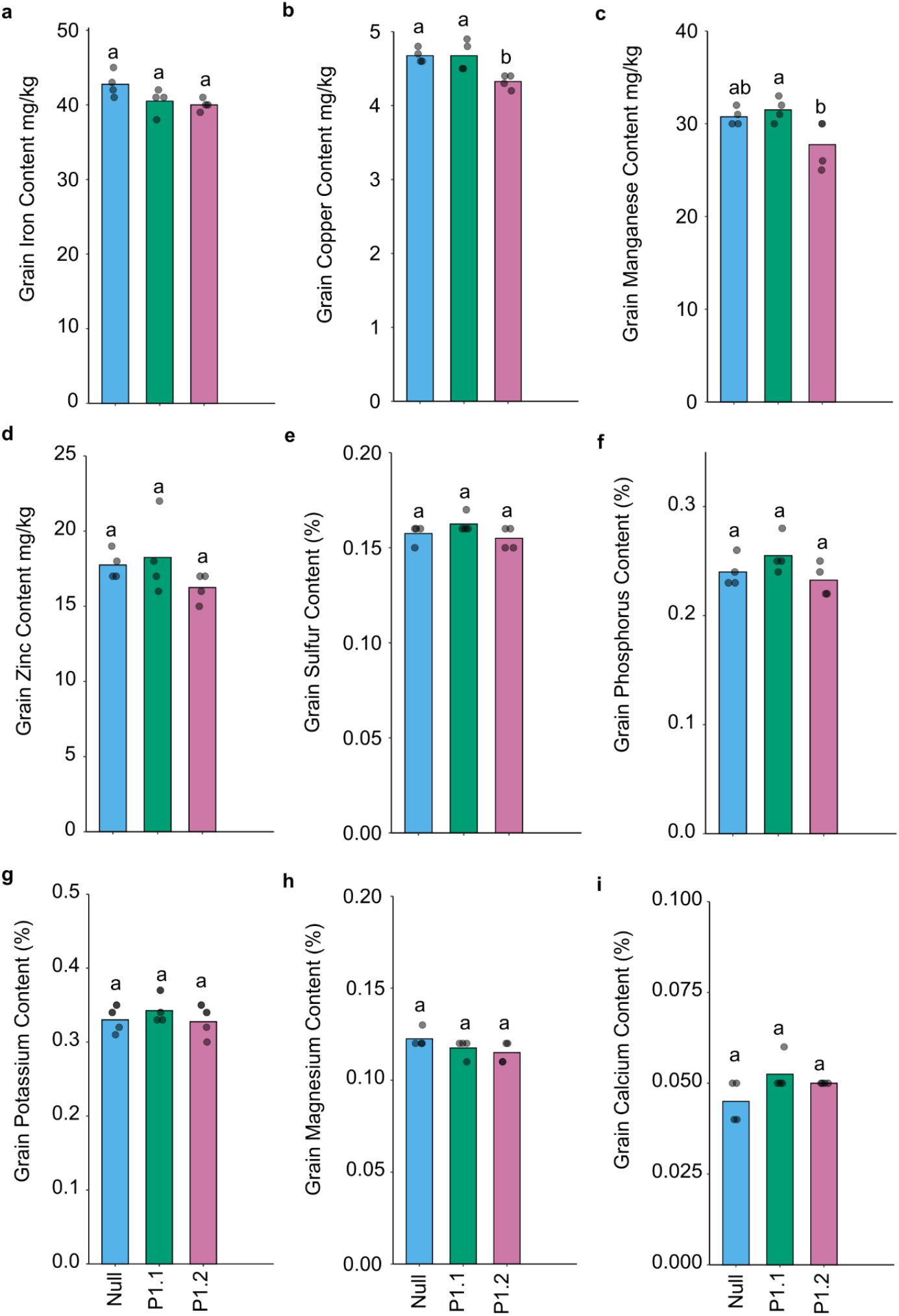
Grain nutrient content of *pUbi::OsPSTOL1* transgenics and their null segregant grown in the field. Characteristics of the grain of wheat cultivated in a field trial at Glenthorne farm in South Australia under nutrient replete conditions. **a-i**, Grain iron (**a**), copper (**b**), manganese (**c**), zinc (**d**), sulfur (**e**), phosphorus (**f**), potassium (**g**), magnesium (**h**), and calcium (**i**) content of *pUbi::OsPSTOL1* transgenic lines and their null segregants (*n*=4). Grain from several plants from each genotype was pooled into four groups for nutrient determination. Means significantly different between genotypes are indicated by different letters (*P<*0.05, analysis of variance (ANOVA) with Tukey’s honest significant difference (HSD) test).

**Figure S10.**
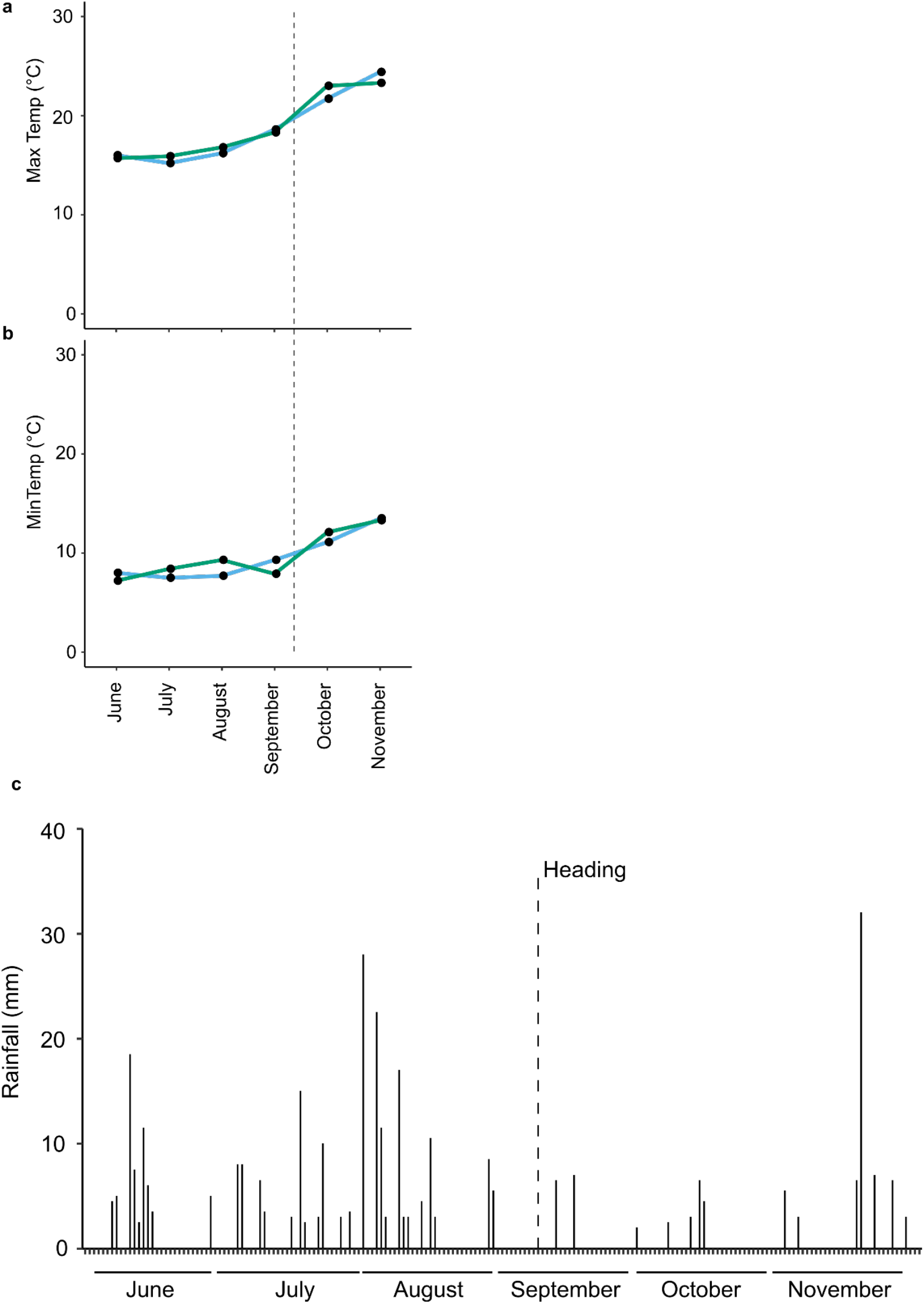
Weather conditions during the field trial. **a-b**, Maximum daily temperature (**a**), minimum daily temperature (**b**), and rainfall per day (**c**) at Glenthorne farm, Australia (−35.056923 S 138.556224 E) where field trials were conducted in 2018. The dashed vertical line indicates the point where the first plants began heading. The 2018 ainfall data was collected directly at Glenthorne farm. All other data was retrieved from the weather station at Adelaide Airport, South Australia. Related to Figure 5.

**Figure S11.**
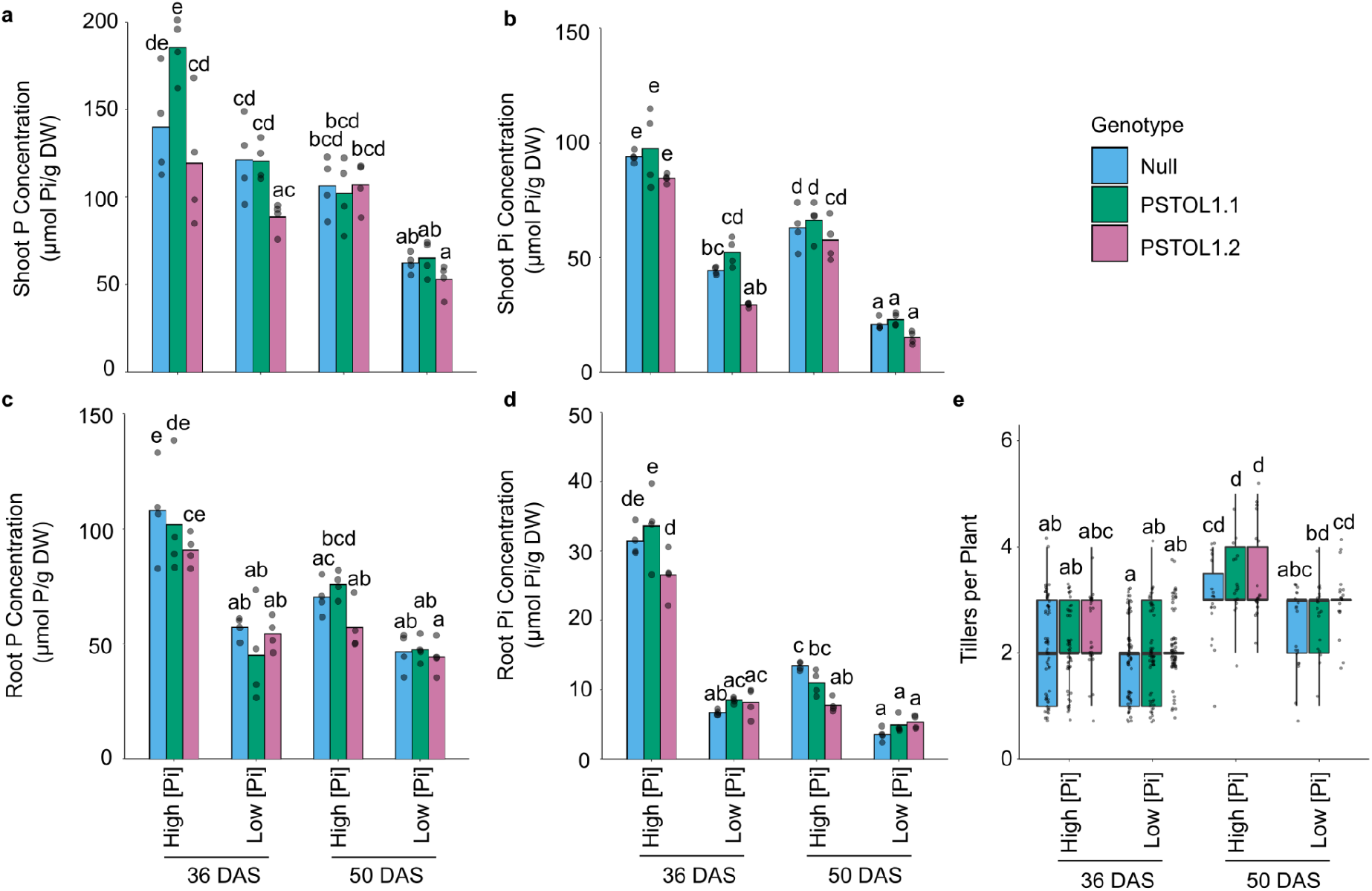
Phosphate status and growth of shoots and roots *pUbi::OsPSTOL1* lines and their null segregant under high and low [Pi] conditions. **a-c**, Total P (**a,c**) and Pi (**b,d**) concentration of shoots (**a,b**) and roots (**c,d**) at 36 and 50 DAS under High and Low [Pi] conditions, n=4. **e**, Number of tillers per plant at 36 and 50 DAS under High and Low [Pi] conditions. 36 DAS, n=60. 50 DAS, n=20. Means significantly different between genotypes are indicated by different letters (*P<*0.05, analysis of variance (ANOVA) with Tukey’s honest significant difference (HSD) test). Supporting data for Figure 6.

**Figure S12.**
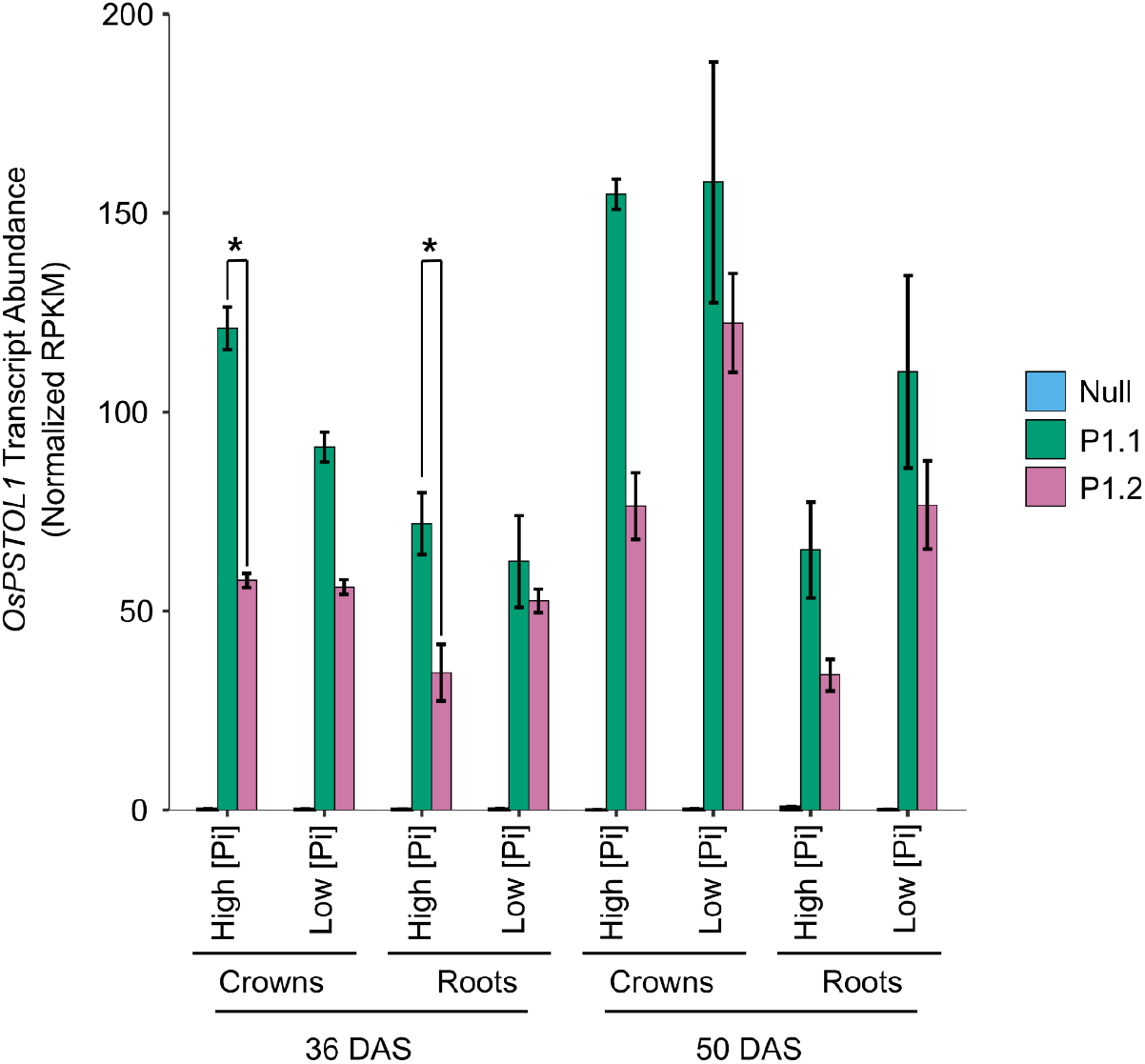
Transcript abundance of the *OsPSTOL1* transgene in wheat p*Ubi*::*OsPSTOL1* transgenic lines. Transcript abundance of *OsPSTOL1* transgene determined by mRNA-seq. Bars indicate standard error. *n*=3. Asterisk indicates significantly different at *P* < 0.05. As would be expected, Null samples that lack *OsPSTOL1* had < 0.25 RPKM mapping to the *OsPSTOL1* transgene sequence added to the wheat genome for read mapping. RPKM values were normalized across the transcriptome.

**Figure S13.**
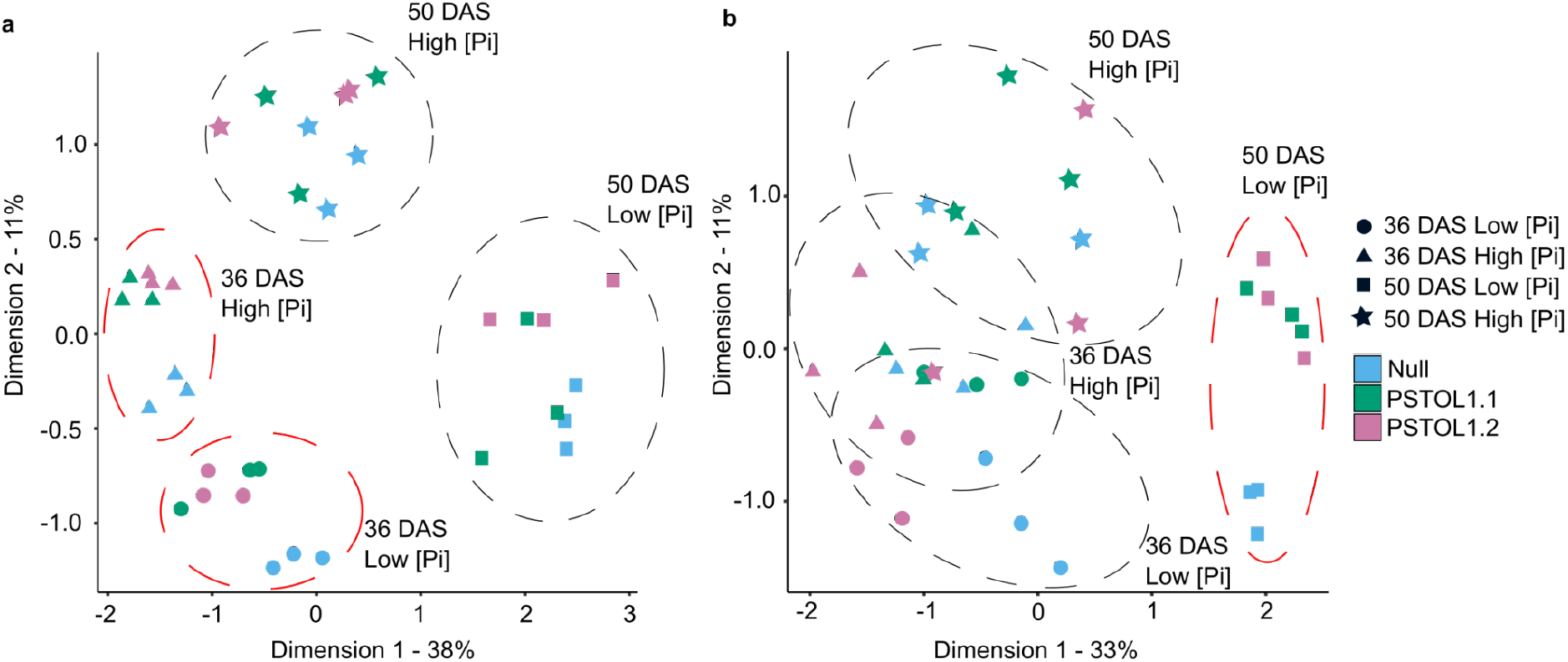
Root and crown transcriptomes of independent *pUbi::OsPSTOL1* transgenics events and their null segregant are influenced by P status and age. **a, b**, Multidimensional scaling plots of transcriptomes of roots (**a**) and crowns (**b**). Groupings indicated with dashed lines. Red groupings with clear separation of genotypes.

**Figure S14.**
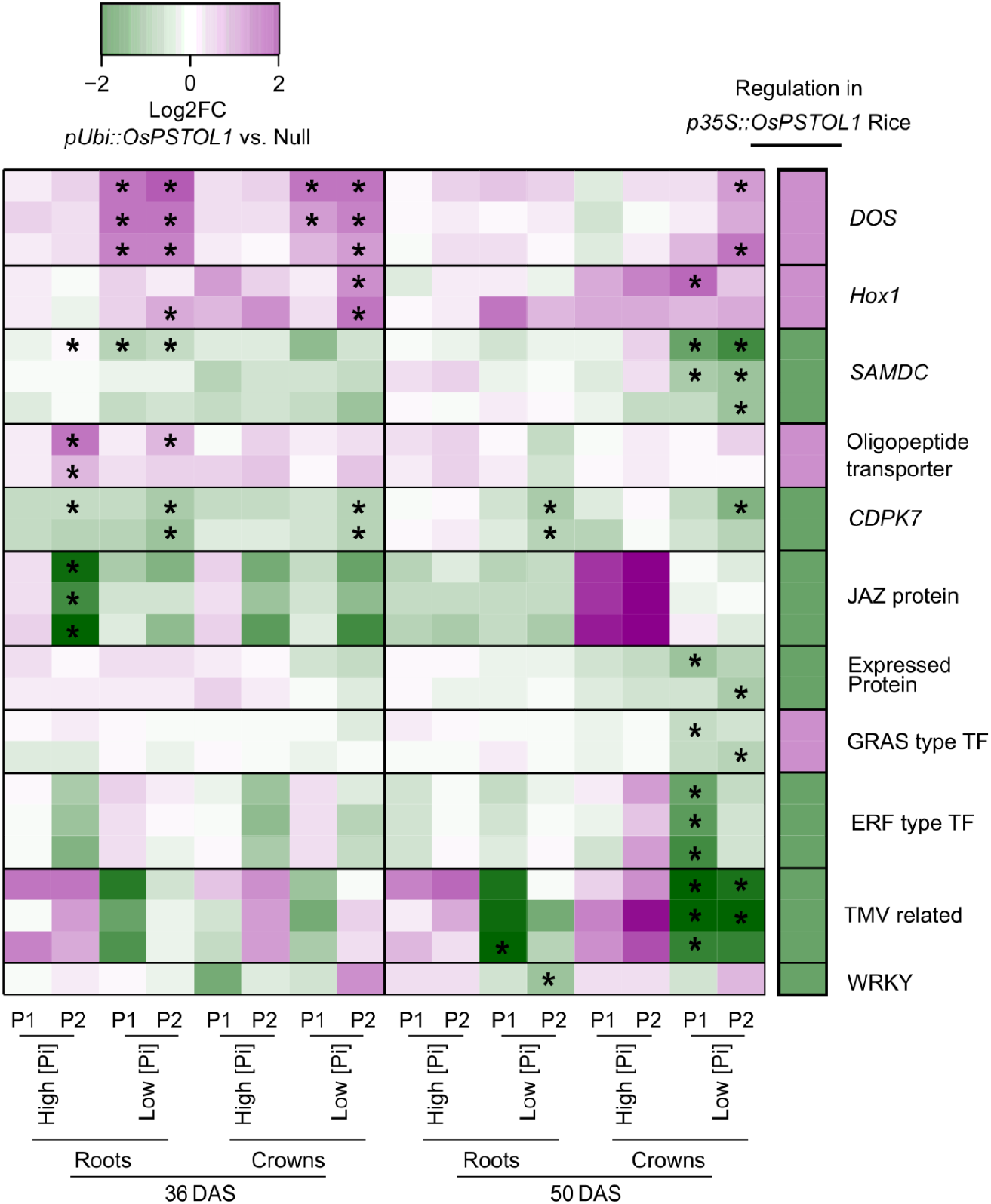
Commonly differentially regulated genes in *OsPSTOL1* transgenics of wheat and rice. Meta-analysis of wheat orthologs of rice genes identified to be constitutively differentially regulated independent of growth condition in roots of heading stage and mid-tillering stage *35S::OsPSTOL1* transgenics (cultivar IR64) relative to non-transgenic nulls (Gamuyao *et al*. 2012). The direction of regulation of each gene by *OsPSTOL1* in the *35S::OsPSTOL1* transgenic rice lines, is indicated in the heatmap on the right. Only wheat homologs of the rice orthologs that were differentially regulated in at least one condition are shown (*P*<0.05) in the heatmap on the left. Significant differential regulation of wheat genes indicated with an *. Rice genes had similar regulation regardless of P condition or growth stage tested (Gamuyao *et al*. 2012).

**Figure S15.**
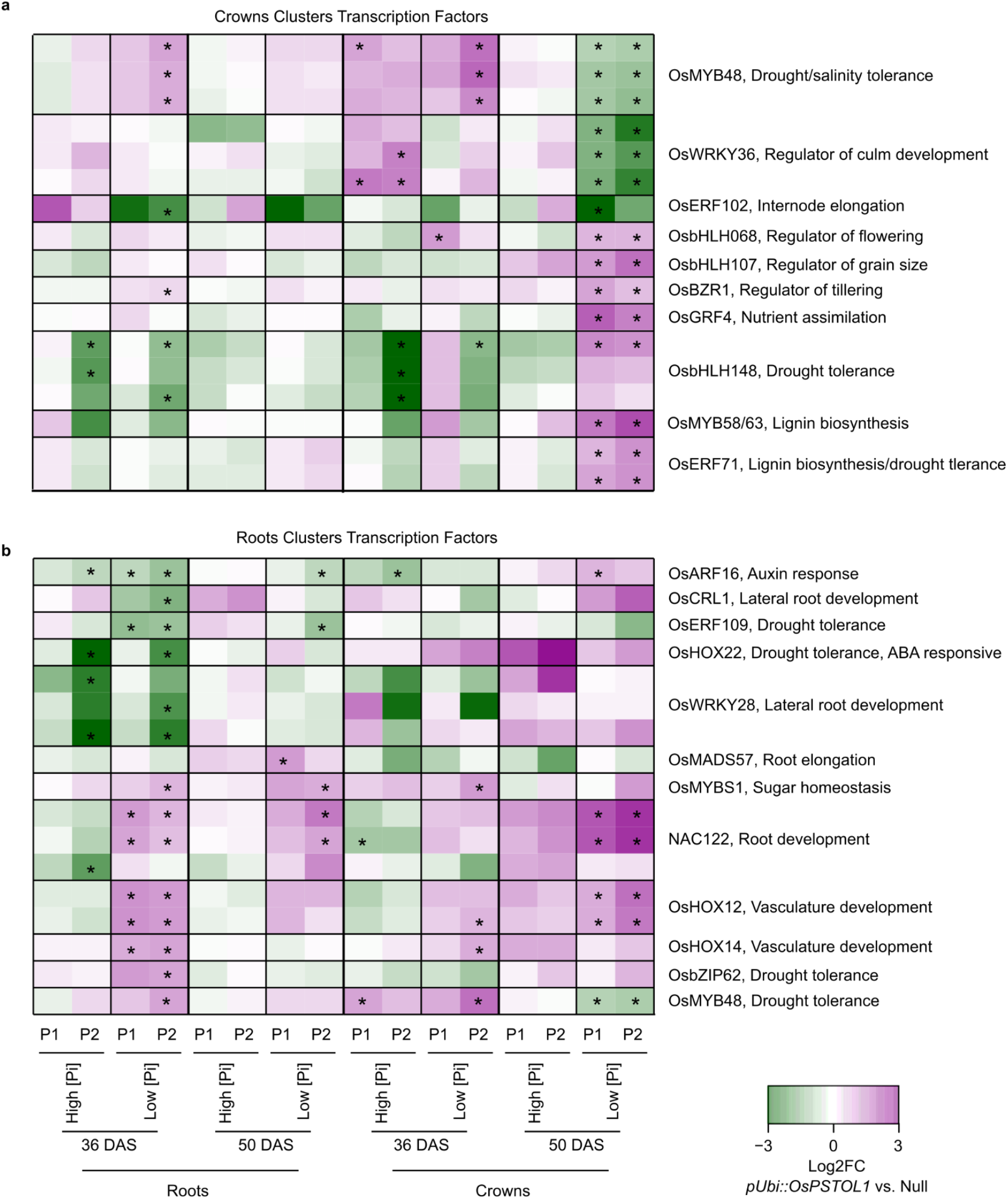
Heatmaps of notable transcription factors differentially regulated in roots and crowns of *pUbi::OsPSTOL1* lines. **a,b** Select transcription factors in crowns (**a**) and roots (**b**) from the clustering analysis in Figure 6e,f. Names and annotations are for the rice ortholog. Wheat orthologs differentially regulated in at least one genotype or condition in *pUbi::OsPSTOL1* transgenic lines compared to null segregants (*P*<0.05) are indicated with an asterisk.

**Figure S16.**
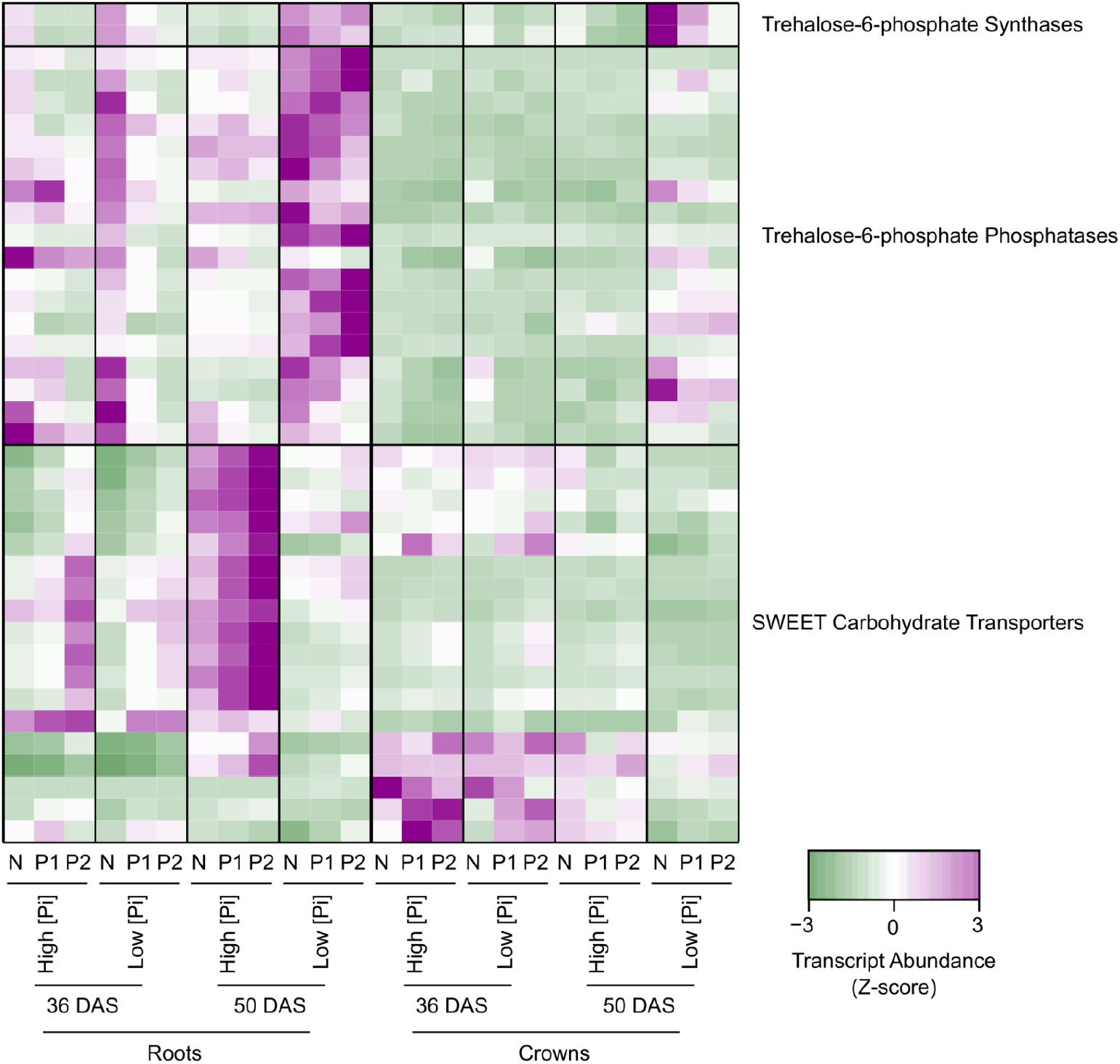
Genes related to carbohydrate allocation are differentially regulated in *pUbi::OsPSTOL1* lines. Heatmap of Z-scores of *TPP*s, *TPS*, and *SWEET* transporters that are involved in allocation and movement of carbohydrates. Genes shown are differentially regulated in at least one transgenic line in at least one condition. Wheat orthologs differentially regulated in *pUbi::OsPSTOL1* transgenic lines compared to null segregants (*P*<0.05) are indicated with an asterisk.

**Figure S17.**
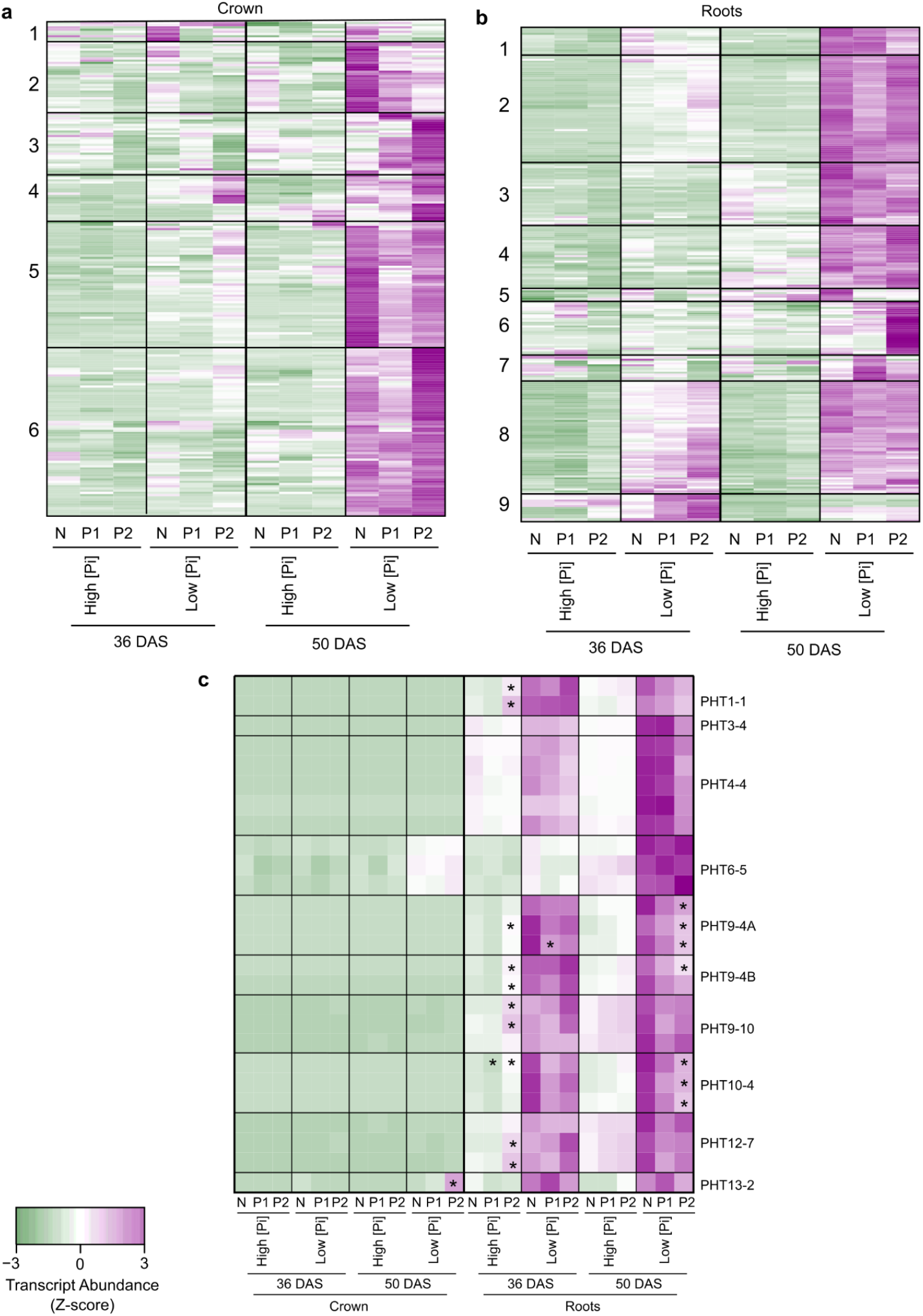
Low P-response marker genes are differentially expressed in the *pUbi::OsPSTOL1* transgenic lines relative to the Null in crowns and roots. **a,b** Cluster analysis of low P marker genes at 36 and 50 DAS under high and low [Pi] in the crowns (**a**) and roots (**b**). Marker genes are wheat orthologs of rice low P signaling and mobilization (). We identified 267 root and 223 crown orthologs expressed in roots or crowns at one or more timepoint, respectively. **c**, Trasnscript abundance of known P transporters. * indicates significantly different than the null segregant at the same time, tissue, and P status (|log2FC| ≥1, *P*<0.05). Genes shown are differentially regulated in at least one transgenic in at least one condition.

## Supporting Materials and Methods

### Determination of genotypic, geographic, and cultivation ecosystem information of rice varieties

DNA sequence information for varieties from the 3000 Rice Genome database was obtained from the Rice Pan-Genome Browser (Sun et al. 2017). A BLASTn search (Altschul et al. 1997) with *OsPSTOL1* (GenBank Accession AB458444.1) of the rice pan-genome identified the J and K-alleles annotated as two distinct loci. The J-allele coding sequence (CDS) is listed as Un_maker_67132. The CDS of the K-allele is unannotated, but another gene within 100 bp of exon 1 (Un_maker_63729) was used as an initial proxy for the K-allele genotype. The presence/absence tool in the Rice Pan-Genome Browser was used to determine the presence or absence of either allele. Varieties lacking either allele were scored as the Absent haplotype. To validate genotype, variant call format (.vcf) files of each variety (Wang et al. 2018) derived from alignments to the *aus* landrace Kasalath genome were examined, where *OsPSTOL1* is present at UM:6007176-6008150. Two known single nucleotide polymorphisms (SNPs) were found at UM6007585 and UM6007586, and designated Kstop1 and Kstop2, respectively. Minor variants of major presence alleles, without nonsense mutations, were not given a separate allele designation. Varieties scored by the browser as the absent haplotype but had variant information across >75% of the CDS were treated as present, whereas any varieties scored as present that had variant information across < 75% of the CDS were subsequently scored as absent haplotype. Varieties with coverage ≥75% of the CDS were marked as present, so as not to discount varieties with low read coverage but enough meaningful SNPs to distinguish the allele. Overall accuracy from the proxy score to browser confirmation was >97%. Varieties with high sequence coverage buy low sequence calls on meaningful SNPs were marked as ambiguous and discarded.

For varieties not in the 3000 Rice Genome dataset, the allele of *OsPSTOL1* was determined by aligning genomic sequence files of landraces to the K-allele of *OsPSTOL1* using Bowtie 2 (Langmead and Salzberg 2012). After the alignment, the consensus sequence was determined from the generated BAM file using the mpileup function in samtools followed by the seqtk package (Li 2016) to generate a FASTQ file of the consensus(Li et al. 2009). BLASTn was used to align the generated consensus to both the J- and K-allele and the allele was determined. Data on allelic variation for *OsPUPk20-2* in varieties in the 3000 Rice Genome database were obtained from RiceVarMap v2.0 using the Haplotype Network Analysis tool (http://ricevarmap.ncpgr.cn/v2/hap_net). Alleles of *OsPUPK20-2* and *OsPUPK67-1* were determined as described above for *OsPSTOL1* using sequences from Nipponbare.

Country-of-origin, subspecies, rice growing ecosystem, and SNP group information was mined from the Rice Pan-Genome Browser (Sun et al. 2017) and (Gutaker et al. 2020). Countries with fewer than 10 varieties sampled were discounted. Regional data on rice growing ecosystem distribution were from (Haefele, Nelson, and Hijmans 2014). P fertilizer application data were from (Roser and Ritchie 2013), updated in 2017. For figure 1f and 3a, andraces from Hong Kong were grouped with China. Landraces from North and South Korea were grouped together. Countries with fewer than 10 landraces were excluded.

### *OsPSTOL1* homologs, phylogenetic tree construction, and protein domain prediction

*OsPSTOL1* orthologs have been reported in some *Oryza(Neelam et al. 2017; Pariasca-Tanaka et al. 2014)*. To identify *OsPSTOL1*-like genes in other species, BLASTP was performed with the K-allele of OsPSTOL1 as a query against reference proteomes of *Arabidopsis* (GCA_000001735.2), sorghum (GCA_000003195.3), maize (GCA_000005005.6), rice (GCA_001433935.1), and wheat (GCA_900519105.1) obtained from UniProt (UniProt Consortium 2021). The top ten hits with E-values < 1e^-100^ from each species were aligned using MAFFT (Katoh and Standley 2013). Gaps were trimmed with trimAl (Capella-Gutiérrez, Silla-Martínez, and Gabaldón 2009) using default settings. These alignments comprise the kinase domain. A draft maximum alignment tree was constructed using FastTree(Price, Dehal, and Arkin 2010). Sequences were manually curated, removing distant outgroups. *Arabidopsis* proteins formed a subgroup of which two representatives were retained. The curated list was realigned, trimmed and a final tree was constructed using RAxML (Kozlov et al. 2019), with default settings and 1000 bootstraps. Nodes with bootstrap values <40 were collapsed with TreeCollapserCL4 (http://emmahodcroft.com/TreeCollapseCL.html). Protein domains were predicted using hmmscan (Finn, Clements, and Eddy 2011), ScanProsite (de Castro et al. 2006), and Phobius (Käll, Krogh, and Sonnhammer 2007).

### Confirmation of transgene presence and expression

Plant DNA was extracted as described (Regmi et al. 2020) and total RNA was extracted using TRIzol reagent (Invitrogen) and a Direct-zol RNA MiniPrep Kit (Zymo Research, CA, USA) with an on-column DNase treatment. *OsPSTOL1* and *TaGAPDH* or *TaACTIN* genes were assayed by PCR amplification with gene specific primers (Table S1a) using the OneTaq® 2× master mix (New England Biolabs Inc., MA, USA). For cDNA synthesis, 2.0 μg of RNA was processed in a High-Capacity cDNA Reverse Transcription Kit (Applied Biosystems, Victoria, Australia) reaction using random oligo primers followed by qRT-PCR (Reynoso et al. 2019), with results normalized to *TaACTIN* in four biological and four technical replicates. Transgene DNA or RNA was checked in 5 seedlings of each independent line in every experiment. Seed of any plant that did not show presence or expression of the transgene in 100% of progeny sampled was discarded.

### Evaluation of transgenic wheat under controlled phosphate nutrition

For evaluation of growth under low P conditions in the greenhouse, seeds were germinated as described above, except presterilized prior to pre-germination on wet filter paper in sealed petri dishes in a chamber at 22°C, with a 16 h light (115 μE) and 8 h dark cycle for 3 d. Five to six germinated seedlings of average size and growth were transplanted into each pot containing a mix of 70% (v/v) #20 silica sand (Gillibrand) and 30% (v/v) Green Grades Profile™. Profile™ contains 10% (v/v) aluminum oxide which acts as a solid-state buffer for P (Lynch et al. 1990). Prior to seedling transplantation, the growth medium of each pot was pre-washed 16 times with 0.75 L Industrial H_2_O, with at least 4 hours between each wash to reduce the starting P content to 25 μM. For the first 14 d after sowing (DAS), plants were sub-irrigated (from pot base) with industrial water containing negligible amounts of P and nitrogen. At 14 DAS, pots were sub-irrigated with 3 L of industrial water supplemented with the additional nutrients. For high [Pi] (HP), the solution contained 436 μM KH_2_PO_4_, and for low [Pi] (LP) this was substituted with 440 μM KCl (Dataset S1c). The industrial water used contains 1625 μM Ca^2+^, 375 μM Mg^2+^, and 656 μM SO_4_^2-^. Trays were refilled daily, and fertilizer solution was changed every 3 d. Plants were harvested at 36 and 50 DAS, representing the mid tillering (Zadoks stage 23) and mid elongation (Zadoks stage 31), respectively. At harvest, roots were quickly photographed and weighed. Roots were then separated from the crown (0.25 cm above the highest, emerged crown root) and the shoot. Each pot (5-6 plants) was pooled as a biological replicate, with four biological replicates in total. These were frozen in liquid nitrogen and cryopreserved at −80°C.

For evaluation of root systems, pre-germinated seeds were transplanted into 40 cm tall, 10.16 cm diameter PVC pipes filled with a mix of 70% (v/v) #20 silica sand (Gillibrand) and 30% (v/v) Green Grades Profile™ to 3 cm below the top. For LP, the medium was washed through with 2 L of water every 4 hours until the P concentration was 5 μM. After transplantation, plants were drip irrigated daily. For HP, the solution contained 436 μM KH_2_PO_4_. For LP, 5 μM KH_2_PO_4_ and 335 μM KCl was used. For the first 5 d after seeding the top of the pipe was covered with a transparent plastic cup to prevent drying. At 28 DAS, root systems were recovered by submerging the pipe in water to gently remove the media. Roots were stored in 50% (v/v) ethanol at 4°C. Root architecture was visualized by suspending the roots in a plastic tray filled with 3 L of water. Tweezers were used to gently separate the roots for photography. Crown roots were counted, residual media was removed, and whole roots were dried at 60°C for at least 72 hours before weighing.

### Identification of differentially regulated genes

Differentially regulated gene (DRG) transcripts were identified by use of the limma-voom package(Law et al. 2014). Raw RNA-seq read counts were normalized with voom using the quantile method. Genes with less than 5 counts per million (CPM) in at least three samples were excluded. After normalization, count data were used to calculate reads per kilobase per million reads (RPKM). The functions lmfit, contrasts.fit, and ebayes were used to calculate differential expression for the varied contrasts, to determine Log_2_ Fold Change (Log_2_FC) and adjusted P values (adj.P.Val). The fdr method was used for controlling the false discovery rate (FDR). Clustering was performed with the Clust package(Abu-Jamous and Kelly 2018) using Z-score normalization of Log_2_FC values. Gene Ontology (GO) enrichment was performed with systemPipeR(H Backman and Girke 2016) using the wheat GO definitions from BioMart (Smedley et al. 2009) (Ensembl Release 50). Transcription factors were subsetted using the Plant Transcription Factor Database (Jin et al. 2017). Wheat gene orthologs of rice genes were found using BioMart.

## Notes

### Competing Interest Statement

The authors have declared no competing interest.

